# Cortical eigenmode coordinates provide compact subject-level signatures across structural MRI, resting-state fMRI, and EEG

**DOI:** 10.64898/2026.05.25.726064

**Authors:** Hyung G. Park, Thaddeus Tarpey

**Affiliations:** Department of Population Health, NYU Grossman School of Medicine, New York, USA

**Keywords:** cortical eigenmodes, Laplace–Beltrami basis, multimodal neuroimaging, structural MRI, resting-state fMRI, EEG

## Abstract

A practical barrier in multimodal neuroimaging is that structural MRI, fMRI, and EEG are often analyzed in modality-specific spaces or reduced to atlas- and sensor-based summaries, limiting the construction of common, interpretable subject-level brain signatures. We evaluate cortical Laplace–Beltrami eigenmode coordinates as a shared geometry-aligned language for structural MRI (sMRI), resting-state fMRI (rs-fMRI), and EEG. In this framework, sMRI morphometric fields are represented by cortical eigenmode coefficients, rs-fMRI by covariance among eigenmode time-series coefficients, and EEG by mode–frequency–condition summaries.

Using the Max Planck Institute Leipzig Mind–Brain–Body dataset (MPI–LEMON), we compared unimodal eigenmode-coordinate summaries, multimodal cortical eigenmode-coordinate PCA, conventional atlas/sensor-based PCA and ridge representations, and *geometric eigenmode multiview factorization* (GEMF). GEMF is a structured decomposition that preserves the modality-native organization of the data objects while separating shared from modality-specific variation. Eigenmode-coordinate representations yielded compact subject-level signatures with strong external validity for chronological age and a secondary cognitive outcome. Multimodal eigenmode-coordinate PCA was among the strongest-performing approaches, reached high age-prediction performance at moderate dimension, and consistently outperformed conventional low-dimensional PCA. GEMF selected an even lower-dimensional shared representation and remained competitive with the benefit of providing interpretable shared and modality-specific factors.

These findings support cortical eigenmode coordinates as a practical foundation for compact, interpretable, and multimodally aligned subject-level brain signatures.

## 1. Introduction

A central practical challenge in multimodal neuroimaging is that structural MRI, fMRI, and EEG are typically represented in different feature spaces before statistical analysis (Sui et al., 2012; Calhoun and Sui, 2016; Jorge et al., 2014). Structural MRI is often summarized by regional morphometric measures, such as cortical thickness, surface area, curvature, or volume; resting-state fMRI by parcel-level connectivity or covariance summaries; and EEG by sensor- or source-level spectral summaries. These choices are convenient and widely used, but they leave investigators with subject-level features that are not naturally comparable across modalities. They can also obscure cortical spatial organization by replacing continuous cortical fields with parcellated regional averages, sensor summaries, or modality-specific coordinate systems whose spatial scales are not directly aligned. As a result, multimodal integration is often handled through feature-level fusion, feature concatenation, modality-specific dimension reduction, or post hoc linking of latent components, rather than through a shared geometry-aligned coordinate system.

This limitation has motivated a large literature on multimodal fusion. Data-driven fusion methods, including joint independent component analysis (ICA), multimodal canonical correlation analysis, linked ICA, joint-and-individual decompositions, and related multiblock factorization approaches, provide powerful tools for detecting cross-modal co-variation and linked latent sources (Sui et al., 2012; Groves et al., 2011; Klami et al., 2013; Lock et al., 2013; Zhou et al., 2016; Calhoun and Sui, 2016). These approaches have been influential in psychiatric, cognitive, and clinical neuroimaging applications, where multimodal features can reveal subject-level patterns not apparent from any single modality alone (Kucukboyaci et al., 2014; Sui et al., 2023). However, most existing pipelines first reduce each modality to features defined in its own native analysis space. This feature-level strategy makes fusion tractable when modalities differ in dimension, units, and sampling structure, but it does not by itself define a common cortical coordinate system across modalities. Thus, there remains a need for representations that make modalities spatially comparable before fusion, rather than relying only on downstream statistical alignment.

This challenge is especially important when the scientific goal is to construct subject-level brain signatures that are compact, interpretable, and useful beyond a single imaging modality. Investigators increasingly need brain feature representations that can support multimodal integration while retaining a clear connection to neurobiology (Arbabshirani et al., 2017; Shen et al., 2017; Sui et al., 2020; Marquand et al., 2019; Rutherford et al., 2022; Ferreira et al., 2022). Atlas- and sensor-based summaries are often effective for specific tasks, such as region-level morphometric association, connectome-based prediction, or sensor-level EEG biomarker analysis, but they do not provide a general representation of cortical organization that is shared across sMRI, fMRI, and EEG and ordered by a biologically meaningful notion of spatial scale.

Cortical Laplace–Beltrami (LB) eigenmodes offer a natural alternative (Seo and Chung, 2011; Pang et al., 2023). Defined on the cortical surface, these eigenmodes form an intrinsic multiscale basis ordered from coarse to fine spatial variation, and can provide a geometry-aligned coordinate system in which different cortical measurements can be represented using a common spatial language. Throughout this paper, we refer to these coordinates as *cortical eigenmode coordinates* or, equivalently, *harmonic coordinates*. The latter terminology emphasizes the analogy with harmonic basis functions on a geometric domain, while the former emphasizes their construction from the cortical LB operator. In either case, the key point is that the coordinates are tied to cortical geometry and ordered by spatial scale.

The central goal of this work is to test whether cortical eigenmode coordinates can serve as a practical foundation for multimodal subject-level brain signatures. We consider two related tasks. First, we seek low-dimensional subject representations that are externally informative and can be compared across modalities. Second, we seek to characterize how shared multimodal variation is distributed from coarse to fine cortical spatial scales.

The LB eigenbasis is well suited to these goals because it provides both a common cortical coordinate system and an explicit multiscale ordering.

Within this common coordinate system, we represent each modality in a form that preserves its native structure. Structural MRI is represented as a matrix of LB-basis coefficients across multiple morphometric fields (Seo and Chung, 2011; Pang et al., 2023; Cao et al., 2024). Resting-state fMRI is represented through covariance among LB-basis time-series coefficients, yielding a subject-specific harmonic covariance object related to prior work on harmonic representations of brain activity (Atasoy et al., 2016; Luppi et al., 2023; Pang et al., 2023). EEG is represented as a mode-by-frequency-by-condition tensor of spectral summaries, motivated by prior work using harmonic or modal decompositions for EEG and large-scale brain dynamics and by recent tensor-based EEG analyses (Wingeier et al., 2001; Nunez and Srinivasan, 2006; Gholamipourbarogh et al., 2024). The framework is not limited to these three modalities in principle: any cortical measurement that can be expressed as a surface field, time-varying surface field, covariance object, or structured tensor over cortical eigenmodes could be incorporated. In this paper, we focus on sMRI, rs-fMRI, and EEG because they provide a practically important test case spanning anatomy, hemodynamics, and electrophysiology.

We evaluate this framework in two complementary ways. First, we ask whether cortical eigenmode-coordinate representations produce compact subject-level signatures with strong external validity. To do this, we compare unimodal eigenmode-coordinate PCA summaries for sMRI, rs-fMRI, and EEG; multimodal eigenmode-coordinate PCA; conventional atlas/sensor-based PCA; and high-dimensional conventional ridge regression in the Max Planck Institute Leipzig Mind–Brain–Body dataset (MPI–LEMON) (Babayan et al., 2019). This PCA-based fusion benchmark is motivated by prior work showing that PCA and related multivariate approaches can integrate multimodal neuroimaging summaries for brain–behavior association (Kucukboyaci et al., 2014; Sui et al., 2023), while here the PCA is applied after all modalities have been expressed in a shared cortical eigenmode coordinate system. Second, we develop and evaluate the proposed geometric eigenmode multiview factorization (GEMF), a structured low-rank decomposition that separates subject-level variation into shared and modality-specific components while preserving the modality-native organization of the data objects.

GEMF is related to a broad literature on multiview and multimodal factorization. Existing approaches include joint ICA and related multimodal fusion models for neuroimaging (Sui et al., 2012, 2013; Groves et al., 2011; Calhoun and Sui, 2016), Bayesian canonical correlation analysis and group factor analysis (Klami et al., 2013, 2015; Ferreira et al., 2022), joint-and-individual variation decompositions such as JIVE (Lock et al., 2013), group component analysis for multiblock data (Zhou et al., 2016), and more general latent factor models for multi-view and multi-omics data, including multi-omics factor analysis (MOFA) (Argelaguet et al., 2018, 2020). These methods provide important tools for identifying shared, linked, and view-specific sources of variation. GEMF differs in two ways that are central to the present application. First, all views are expressed in a common cortical eigenmode coordinate system, so shared factors are interpretable with respect to cortical spatial scale. Second, GEMF preserves the native structure of each modality rather than reducing all inputs to a single vector form: sMRI is modeled as a matrix over harmonic mode and morphometric feature, rs-fMRI as a symmetric harmonic covariance matrix, and EEG as a mode–frequency–condition tensor. This structure-preserving formulation is especially relevant for tensor-valued neurophysiological data, where vectorization can obscure mode-specific organization (Bi et al., 2021; Gholamipourbarogh et al., 2024). Thus, GEMF estimates shared latent subject factors jointly across sMRI, rs-fMRI, and EEG, while modality-specific factors capture residual structure unique to each measurement type.

Our empirical results support three main conclusions. First, cortical eigenmode coordinates yield compact subject-level signatures with strong external validity for chronological age and a secondary cognitive outcome. Second, simple multimodal fusion in this common cortical harmonic language is effective: multimodal harmonic-coordinate PCA performs strongly, reaches high age-prediction performance at moderate dimension, and consistently outperforms conventional low-dimensional PCA. Third, GEMF provides a compact and interpretable multiview decomposition: it selects a low-dimensional shared representation, remains competitive in prediction, and enables visualization of shared and modality-specific structure across cortical scale, fMRI harmonic-coordinate covariance, and EEG frequency/condition profiles. Together, these results suggest that the value of cortical eigenmode coordinates is not only dimensionality reduction, but also the ability to define compact, interpretable, and multimodally aligned subject-level brain signatures in a common cortical language.

## 2. Methods

### 2.1. Overview

The main goal of this paper is to evaluate whether cortical harmonic coordinates provide compact, interpretable, and externally valid subject-level brain signatures across multiple modalities, namely structural MRI (sMRI), resting-state fMRI (rs-fMRI), and EEG. Our emphasis is on the representation itself: whether expressing multiple modalities in a common Laplace–Beltrami (LB) coordinate system yields subject-level summaries that are more informative than conventional atlas- or sensor-based summaries and that support feasible multimodal fusion.

The analyses are organized around three questions. First, do harmonic-coordinate representations produce compact subject-level signatures with strong external validity in individual modalities? Second, does a simple multimodal harmonic PCA representation preserve or improve predictive utility relative to conventional atlas/sensor-based multimodal representations? Third, can a structured multiview decomposition separate shared variation from modality-specific variation while retaining competitive subject-level information and providing additional interpretability? We address these questions in the MPI–LEMON cohort using chronological age as the primary external validation target and Leistungsprüfsystem subtest 1 (LPS-1), a cognitive performance measure included in MPI–LEMON, as a secondary cognitive outcome (Horn, 1983; Babayan et al., 2019). A key feature of the framework is that each modality is first represented in a common cortical harmonic coordinate system while preserving its native data structure. Structural MRI is represented as a matrix of harmonic coefficients across multiple cortical morphometric fields. Resting-state fMRI is represented through covariance among harmonic coefficient time series. EEG is represented as a mode-by-frequency-by-condition tensor of resting-state spectral summaries. From these modality-native objects, we construct several subject-level representations: unimodal harmonic-coordinate PCA summaries (Section 2.4.1), multimodal harmonic-coordinate PCA (Section 2.4.2), conventional atlas/sensor-based representations (Section 2.4.3), and geometric eigenmode multiview factorization (GEMF), a structured low-rank decomposition that separates shared and modality-specific variation (Section 2.5).

### 2.2. Common cortical eigenbasis

Let (ℳ, *g*) denote the cortical surface as a two-dimensional Riemannian manifold embedded in three-dimensional space, where *g* is the metric induced by the cortical surface geometry. In computation, ℳis represented by a triangulated cortical surface mesh. We define the first *K* eigenfunctions 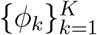 of the Laplace–Beltrami operator Δ_ℳ_ by

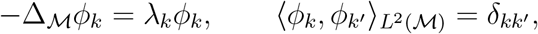

with eigenvalues ordered as

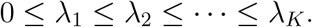

The eigenfunctions form an orthonormal basis for square-integrable functions on the cortical surface. We refer to this basis as the *LB eigenbasis* or the *cortical harmonic basis* (Reuter et al., 2009; Seo and Chung, 2011; Pang et al., 2023). Lower-order basis functions correspond to coarse, long-wavelength spatial patterns, whereas higher-order basis functions represent progressively finer spatial variation. This ordering by spatial scale is important to the present work because it provides both a common coordinate system across modalities and a biologically interpretable multiscale axis.

### 2.3. Modality-native harmonic data objects

#### 2.3.1. Structural MRI

For subject *i* = 1, …, *n*, let *m*_*ij*_(*x*) denote the *j*th cortical morphometric field evaluated at cortical location *x* ∈ ℳ, where *j* = 1, …, *q* and *q* is the number of morphometric features. In the present application, these morphometric features include cortical thickness, sulcal depth, mean curvature, and surface area. Each field is expanded in the common eigenbasis as

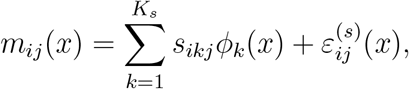

where *K*_*s*_ is the number of structural harmonic modes retained for the sMRI representation. This yields the subject-specific coefficient matrix 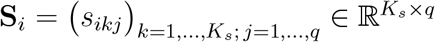. Because the morphometric fields are measured on different numerical scales, coefficients are standardized across features by defining

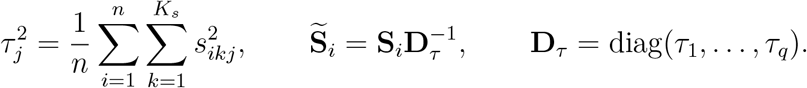

For the PCA-based MPI–LEMON benchmark analyses described in Section 4.1, the full harmonic sMRI feature block retained *K*_*s*_ = 50 modes, providing a low-dimensional summary of coarse-to-intermediate cortical morphometric variation.

#### 2.3.2. Resting-state fMRI

Let *y*_*i*_(*x, t*) denote the cortical rs-fMRI signal for subject *i* at cortical location *x* ∈ ℳ and time *t*. We expand each time-varying spatial field in the same cortical eigenbasis:

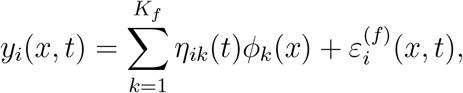

where *K*_*f*_ is the number of harmonic modes retained for the functional MRI representation, and *η*_*ik*_(*t*) is the harmonic coefficient time series for mode *k*. Let 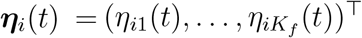. We summarize subject-level rs-fMRI organization by the covariance matrix of these harmonic coefficient time series:

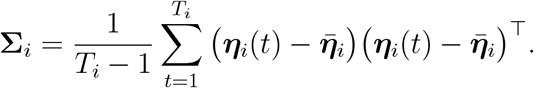

The diagonal entries of **∑**_*i*_ quantify mode-wise temporal variance, while off-diagonal entries summarize cross-mode coupling. For modeling and feature construction, we use the log-Euclidean transform

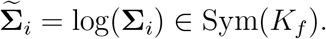

The full harmonic rs-fMRI representation used in the PCA-based benchmark analyses retained *K*_*f*_ = 50 modes, providing a low-dimensional summary of coarse-to-intermediate spatial patterns of functional variance and cross-mode coupling.

#### 2.3.3. EEG

EEG was represented in the same cortical harmonic coordinate system using a forward-projected cortical harmonic dictionary. Let **L** ∈ ℝ^*M* ×*V*^ denote the template EEG leadfield matrix mapping cortical source amplitudes at *V* surface vertices to *M* scalp sensors, and let 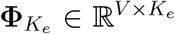 denote the first *K*_*e*_ cortical harmonic modes evaluated on the cortical surface. The forward-projected harmonic dictionary is

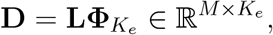

so that each column of **D** is the predicted scalp topography of one cortical harmonic mode. Here *M* is the number of EEG sensors and *K*_*e*_ is the number of retained EEG harmonic modes (Nunez and Srinivasan, 2006; Michel and Brunet, 2019; Park, 2026). In MPI–LEMON, resting EEG was recorded under two conditions, eyes-open and eyes-closed rest. For subject *i*, frequency bin *f*, and condition *c* ∈ {EO, EC}, let

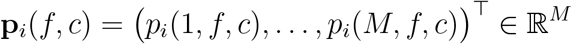

denote the sensor-space spectral-power topography, i.e., the vector of spectral power values across sensors at frequency bin *f* and condition *c* (Nunez and Srinivasan, 2006; Cohen, 2014). We work with log-power topographies,

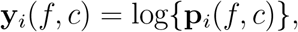

where the logarithm is applied elementwise. The log-power topography is then approximated in the forward-projected harmonic dictionary **D** as

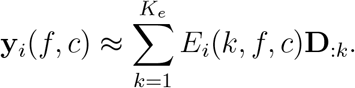

Thus, *E*_*i*_(*k, f, c*) denotes the harmonic coefficient of the log EEG spectral-power topography for mode *k*, frequency bin *f*, and condition *c*. These coefficients are collected into the tensor

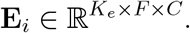

In the present dataset, the EEG representation retained *K*_*e*_ = 20 modes, *F* = 20 frequency bins, and *C* = 2 resting conditions (EO and EC). The smaller number of EEG harmonic modes *K*_*e*_ = 20 reflects the limited spatial bandwidth of scalp EEG and the smoothing induced by volume conduction and the EEG forward operator (Nunez and Srinivasan, 2006; Michel and Brunet, 2019; Park, 2026).

### 2.4. Subject-level representation families

We constructed several subject-level representation families to evaluate whether harmonic coordinates provide useful low-dimensional brain signatures.

#### 2.4.1. Unimodal harmonic PCA

For each modality, the harmonic-coordinate object defined above was converted into a subject-level feature block:

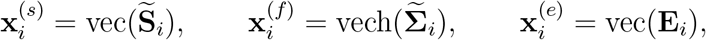

where vech(·) denotes the upper-triangular vectorization of a symmetric matrix. Within each training fold, each feature block was standardized feature-wise and reduced by modality-specific PCA to rank *r*. The same value of *r* was used for each modality-specific PCA representation so that all rank-*r* representations had the same downstream dimension; in the rank-selection analysis, the rank was selected separately within each representation family. The resulting rank-*r* score vectors define the sMRI harmonic-coordinate PCA, fMRI harmonic-coordinate PCA, and EEG harmonic-coordinate PCA representations. These unimodal harmonic-coordinate PCA representations evaluate the subject-level information contained in each modality after projection into cortical harmonic coordinates.

#### 2.4.2. Multimodal harmonic PCA

To evaluate simple multimodal fusion in the common harmonic coordinate system, we concatenated the three harmonic feature blocks:

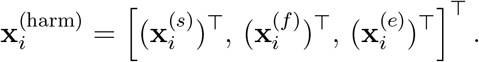

Within each training fold, this concatenated feature vector was standardized and reduced by PCA to rank *r*. The resulting rank-*r* score vector defines the multimodal harmonic-coordinate PCA representation, with representational complexity controlled by the number of retained principal components, *r*. This representation provides a simple and transparent multimodal fusion approach: it uses a standard linear dimension reduction method after all three modalities have been expressed in the same geometry-aligned harmonic coordinate system.

#### 2.4.3. Conventional atlas/sensor-based baselines

We also constructed conventional atlas- and sensor-based comparison features. These features represent commonly used non-harmonic summaries of the same modalities and provide reference representations against which the harmonic-coordinate representations can be evaluated. For sMRI, we extracted Destrieux atlas summaries from FreeSurfer, including regional cortical thickness, sulcal depth, surface area, and mean curvature (Destrieux et al., 2010; Fischl, 2012). For rs-fMRI, we extracted Destrieux surface parcel time series from preprocessed surface-space BOLD data, applied confound regression and band-pass filtering, and summarized each subject by the upper triangle of the parcel correlation matrix, following common atlas-based functional connectivity practice (Biswal et al., 2010; Power et al., 2011). For EEG, we computed conventional sensor-space resting features separately for eyes-open and eyes-closed data, including regional theta, alpha, and beta power summaries and alpha peak frequency (Klimesch, 1999; Nunez and Srinivasan, 2006) (see Supplementary Materials Section S3 for details).

From these conventional feature blocks, we formed two baseline representations. The conventional PCA baseline concatenates all conventional features across sMRI, rs-fMRI and EEG, standardizes them within training folds, and reduces them to rank *r* by PCA. This baseline provides a low-dimensional comparison to multimodal or unimodal harmonic-coordinate PCA representations at the same representation dimension. The conventional ridge regression baseline uses the full standardized conventional feature vector directly in ridge regression. This baseline provides a high-dimensional conventional reference and is not constrained to have the same low-dimensional representation size.

### 2.5. Geometric eigenmode multiview factorization (GEMF)

As a structured multiview representation, we fit geometric eigenmode multiview factorization (GEMF), which decomposes the modality-native harmonic objects into shared and modality-specific components. The harmonic-coordinate PCA analyses used modality-specific harmonic dimensions (*K*_*s*_, *K*_*f*_, *K*_*e*_) = (50, 50, 20). For GEMF, we used a common harmonic dimension *K* = 20 across modalities, obtained by truncating the sMRI and rs-fMRI harmonic objects to the first 20 modes and using the EEG object as constructed above. This focuses the structured multiview decomposition on coarse-to-intermediate cortical organization, which is the spatial range most directly comparable across sMRI, rs-fMRI, and scalp EEG.

Under a fixed-rank model with shared rank *R* and modality-specific ranks (*L*_*s*_, *L*_*f*_, *L*_*e*_), GEMF jointly models the modalities

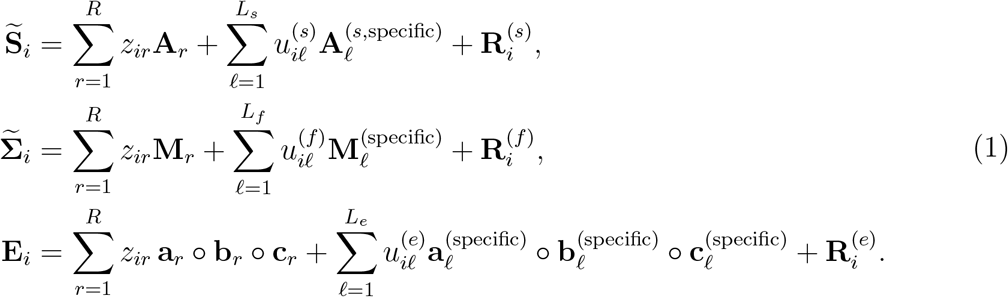

Here, ◦denotes the vector outer product, so that **a**_*r*_◦ **b**_*r*_ ◦**c**_*r*_ is a rank-one tensor over harmonic mode, frequency, and resting-state condition. The vector **z**_*i*_ = (*z*_*i*1_, …, *z*_*iR*_) ∈ ℝ^*R*^ denotes the shared score vector for subject *i* across modalities. We do not impose identical loading patterns for **z**_*i*_ across modalities; rather, each shared score modulates a modality-specific loading object, allowing the same subject-level latent factor to be expressed differently in sMRI, rs-fMRI, and EEG. The modality-specific score vectors are

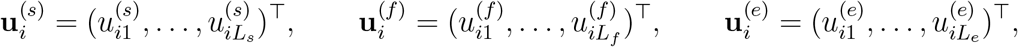

for sMRI, rs-fMRI, and EEG, respectively. The shared components (**z**_*i*_) capture subject-level variation expressed jointly across modalities in the common harmonic coordinate system. The modality-specific components (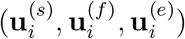) capture structured residual variation unique to each modality.

GEMF preserves the native organization of each modality: sMRI loadings **A**_*r*_ are matrices over harmonic mode and morphometric feature, rs-fMRI loadings **M**_*r*_ are symmetric matrices over pairs of harmonic modes, and EEG loadings **a**_*r*_ ◦ **b**_*r*_ ◦ **c**_*r*_ are rank-one tensors over mode, frequency, and condition. Thus, GEMF provides subject-level shared scores **z**_*i*_ and interpretable loading objects (**A**_*r*_, **M**_*r*_, and **a**_*r*_◦ **b**_*r*_◦ **c**_*r*_) that can be examined across cortical spatial scale, fMRI harmonic covariance structure, and EEG spectral/condition profiles.

#### Penalized objective

GEMF is estimated by minimizing a weighted penalized least-squares objective across modalities:

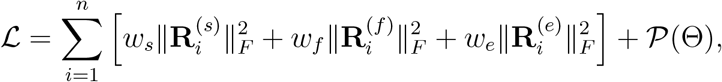

where 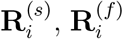 and 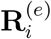 are the residual terms in Equation 1, Θ denotes the collection of loading parameters, and *w*_*s*_, *w*_*f*_, *w*_*e*_ are nonnegative modality weights. In our empirical analyses, we used equal modality weights, *w*_*s*_ = *w*_*f*_ = *w*_*e*_ = 1. The penalty 𝒫(Θ) applies eigenvalue-weighted spatial regularization to shared sMRI, rs-fMRI, and EEG loading components and a smoothness penalty to shared EEG frequency profiles:

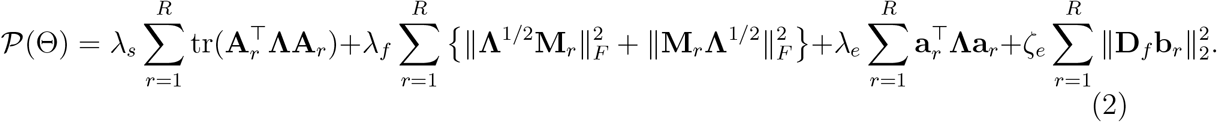

Here **Λ** = diag(*λ*_1_, …, *λ*_*K*_) contains the LB eigenvalues, and **D**_*f*_ is a second-difference matrix over ordered EEG frequency bins, as commonly used for smoothness regularization in penalized spline and finite-difference smoothing methods (Eilers and Marx, 1996; Ruppert et al., 2003). This regularization favors smoother, lower-spatial-frequency shared structure unless higher-order modes are supported by the data. No smoothness penalty is applied to the EEG condition loading **c**_*r*_, because the condition mode contains only the two resting-state conditions and is treated as an unordered condition contrast rather than an ordered axis.

#### Subject-level GEMF representations

We evaluate two subject-level representations derived from GEMF. The first uses only the shared score vector **z**_*i*_ ∈ ℝ^*R*^ and is denoted “GEMF (shared).” This representation has dimension *R* and is directly comparable to a rank-*R* PCA representation. The second uses the concatenated score vector

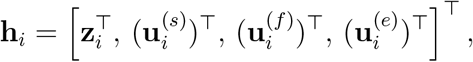

and is denoted “GEMF (full).” When (*L*_*s*_, *L*_*f*_, *L*_*e*_) = (1, 1, 1), GEMF (full) has dimension *R* + 3. This representation allows downstream predictive information to reside in both shared and modality-specific components.

#### Computation

GEMF was fit by block-coordinate minimization of the penalized objective. The unknowns are the shared subject scores **z**_*i*_, the modality-specific scores 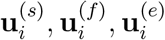,and the corresponding loading objects for sMRI, rs-fMRI, and EEG in Equation 1. Given the current loading objects, the shared and modality-specific subject scores are updated by penalized least squares. Given the current subject scores, the sMRI and rs-fMRI loading objects are updated by penalized matrix regression, while the EEG loading factors are updated by an alternating least-squares step for the mode, frequency, and condition factors (Kolda and Bader, 2009; Sidiropoulos et al., 2017).

The shared loading objects are regularized using the scale-aware penalty in Equation 2. Specifically, the sMRI and EEG spatial-mode loadings (**A**_*r*_ and **a**_*r*_) are penalized by the LB eigenvalues, the rs-fMRI loading matrices (**M**_*r*_) are penalized along both harmonic-mode dimensions, and the EEG frequency profiles (**b**_*r*_) are penalized by a second-difference smoothness penalty. Modality-specific loading objects in Equation 1 are included to absorb structured residual variation unique to each modality; computational details for their weak ridge regularization are provided in Supplementary Materials Section S2. After score updates, score columns are centered and rescaled, followed by rescaling of the associated loading objects, providing a practical identifiability convention. In our application analyses, we fixed (*L*_*s*_, *L*_*f*_, *L*_*e*_) = (1, 1, 1) and varied the shared rank *R*.

Convergence was assessed by the relative change in the penalized objective. Full computational details, including the closed-form score updates and loading-update equations, are provided in Supplementary Materials Section S2.

### 2.6. External validation and rank selection

All predictive comparisons were performed using repeated outer cross-validation. Within each outer training fold, we performed feature standardization, PCA estimation, GEMF fitting, score inference, and rank selection using training data only. Sex was included as an adjustment variable in all prediction models. For continuous outcomes, prediction performance was summarized using the increase in cross-validated *R*^2^ (evaluated on the testing folds) relative to a sex-only baseline,

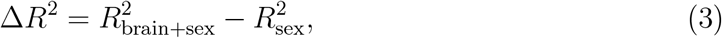

as well as prediction–outcome correlation and root mean squared error (RMSE).

We used two complementary evaluation strategies. First, in the “dimension-matched” comparison, rank-based representations were evaluated over the grid *r* ∈ {1, 2, …, 12}. For PCA-based methods, *r* denotes the number of principal components. For GEMF, *r* denotes the shared rank *R*. In addition, GEMF (full) is shown as a secondary comparison because, for shared rank *R* = *r* and (*L*_*s*_, *L*_*f*_, *L*_*e*_) = (1, 1, 1), its downstream representation dimension is *r* + 3 rather than *r*.

Second, we performed reconstruction-based rank selection using nested cross-validation and the one-standard-error rule (Hastie et al., 2009).Specifically, within each outer training set, candidate ranks were evaluated by an inner cross-validation loop, again using only the training subjects from that split. For each rank-based representation family, candidate ranks were selected from the same grid *r* = 1, …, 12. For a validation fold 𝒱in the inner loop, the normalized reconstruction error (NRE) for a PCA feature block **x**_*i*_ from Section 2.4 was

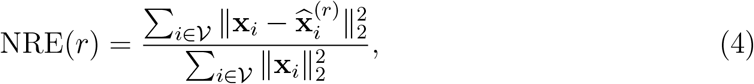

Where 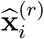 denotes the rank-*r* PCA reconstruction of the corresponding feature block, using PCA estimated from the inner-training fold.

For GEMF, we used a reconstruction criterion based on the three residual terms in the GEMF objective, with each modality normalized by its own total validation-set sum of squares:

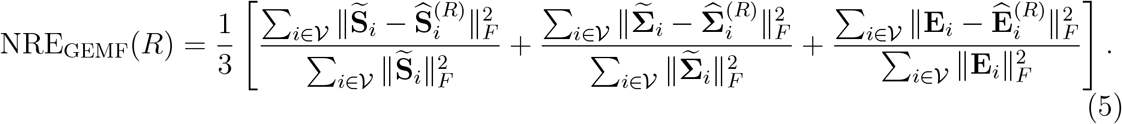

The selected rank was the smallest rank whose mean inner-validation reconstruction error was within one standard error of the minimum. Because rank selection used only reconstruction within the training data, it was independent of the external validation outcomes.

## 3. Simulation study

The simulation study evaluated whether GEMF can recover known shared and modality-specific structure under controlled data-generating mechanisms that match the model class. We generated multimodal data directly in the same object spaces used by GEMF: a standardized sMRI coefficient matrix 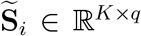, a log-Euclidean rs-fMRI harmonic covariance object 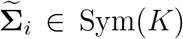, and an EEG mode–frequency–condition tensor **E**_*i*_ ∈ ℝ^*K*×*F* ×*C*^. We fixed (*K, q, F, C*) = (20, 4, 16, 2) and used true shared rank *R* = 3. When modality-specific structure was present, we used (*L*_*s*_, *L*_*f*_, *L*_*e*_) = (1, 1, 1). Sample size was varied over *n* ∈ { 50, 100, 200}, with 100 replications per setting.

For each modality, the simulated object was decomposed into shared, modality-specific, and noise components,

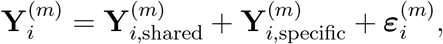

where *m* ∈ {*s, f, e*} denotes sMRI, rs-fMRI, or EEG. The three components were scaled so that their total squared Frobenius norms, summed across subjects, followed prespecified proportions within each modality. We considered three scenarios: 1) a “shared-only” scenario with proportions (0.70, 0.00, 0.30) for shared, modality-specific, and noise components, respectively; 2) a “shared+specific” scenario with proportions (0.45, 0.35, 0.20); and 3) a “weak-specific” scenario with proportions (0.35, 0.05, 0.60). The first scenario evaluates recovery when systematic variation is entirely shared across modalities. The second evaluates recovery when both shared and modality-specific components contribute substantially. The third evaluates whether the shared representation remains stable when modality-specific structure is weak relative to noise.

Recovery was assessed using two primary metrics. Shared-score recovery was measured by the mean canonical correlation between the estimated and true shared score subspaces. Specifically, if 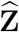 and **Z** denote the estimated and true shared score matrices, we orthonormalized their column spaces and computed the singular values of the resulting cross-product; the reported value is the average of these canonical correlations. Shared-component reconstruction was measured by

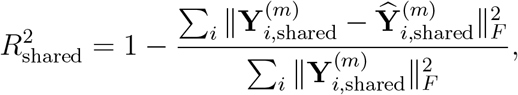

separately for each modality. Modality-specific recovery and rank-selection performance are reported in the Supplementary Materials Section S4.

Table 1 summarizes shared-structure recovery. In the “shared-only” scenario, GEMF recovered the shared score subspace almost perfectly across all sample sizes, and shared-component reconstruction improved with increasing *n*. In the “shared+specific” scenario, recovery was less stable at *n* = 50, particularly for the EEG tensor component, but improved substantially by *n* = 100 and remained strong at *n* = 200. In the “weak-specific” scenario, shared-score recovery remained high and shared-component reconstruction improved with sample size, indicating that the shared representation is stable even when modality-specific structure is weak and noisy. Supplementary analyses in Table S1 showed that modality-specific recovery was strongest in the shared+specific scenario and more variable in the weak-specific scenario. In the supplementary rank-selection experiment in Table S2, selecting the rank by the raw minimum of cross-validated reconstruction error often favored larger ranks, whereas the one-standard-error rule selected ranks close to the true value *R* = 3, with exact recovery most frequent in the shared-only and weak-specific scenarios. Together, these results support GEMF as an interpretable decomposition model whose shared component can be reliably recovered under known data-generating mechanisms.

**Table 1:** Fixed-rank simulation recovery of shared structure across sample sizes. Values are mean (SD) across 100 replications. Shared-score recovery is the mean canonical correlation between the estimated and true shared score subspaces. Shared reconstruction is the 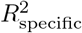 for the true shared component in each modality.

| Scenario | $n$ | Shared-score recovery | Shared $R^2$: sMRI | Shared $R^2$: fMRI | Shared $R^2$: EEG |
| --- | --- | --- | --- | --- | --- |
| Shared-only | 50 | 0.999 (0.000) | 0.952 (0.015) | 0.952 (0.014) | 0.965 (0.058) |
|  | 100 | 0.999 (0.000) | 0.974 (0.009) | 0.974 (0.009) | 0.977 (0.048) |
|  | 200 | 0.999 (0.000) | 0.985 (0.005) | 0.985 (0.005) | 0.986 (0.033) |
| Shared+specific | 50 | 0.939 (0.119) | 0.897 (0.113) | 0.898 (0.110) | 0.751 (0.452) |
|  | 100 | 0.976 (0.079) | 0.953 (0.076) | 0.954 (0.070) | 0.905 (0.287) |
|  | 200 | 0.971 (0.087) | 0.962 (0.073) | 0.961 (0.079) | 0.883 (0.331) |
| Weak-specific | 50 | 0.994 (0.001) | 0.863 (0.017) | 0.866 (0.015) | 0.947 (0.046) |
|  | 100 | 0.994 (0.001) | 0.923 (0.010) | 0.926 (0.010) | 0.955 (0.060) |
|  | 200 | 0.994 (0.000) | 0.955 (0.006) | 0.957 (0.005) | 0.971 (0.039) |

## 4. Application to multimodal subject-level signatures in MPI–LEMON

We evaluated whether multiscale cortical organization, expressed in a common harmonic coordinate system, yields compact subject-level brain signatures with strong external validity. MPI–LEMON provides an appropriate testbed because it includes structural MRI, resting-state fMRI, and resting EEG from the same participants across a broad adult age range, together with demographic and cognitive phenotyping (Babayan et al., 2019). The primary common-cohort analysis used 188 subjects with complete sMRI, rs-fMRI, and eyes-open/eyes-closed EEG harmonic representations after preprocessing and harmonization. Briefly, cortical morphometric fields were projected to the LB basis to form sMRI coefficient matrices; surface-space resting-state BOLD time series were projected to the same basis and summarized by log-Euclidean harmonic covariance matrices; and resting EEG spectral topographies were projected to a forward-projected harmonic dictionary to form mode–frequency–condition tensors. Conventional atlas/sensor-based features were constructed for comparison. Additional preprocessing and feature-construction details are provided in Supplementary Materials Sections S1 and S3. Chronological age was used as the primary external validation outcome, and LPS-1 was used as a secondary cognitive outcome. In all predictive analyses, sex was included as an adjustment variable and performance was summarized primarily by Δ*R*^2^ as defined in Equation 3; prediction–outcome correlation and RMSE were used as additional summaries.

We compared the representation families defined in Section 2.4: unimodal harmonic-coordinate PCA summaries for sMRI, rs-fMRI, and EEG; multimodal harmonic-coordinate PCA; conventional atlas/sensor-based PCA; a high-dimensional conventional ridge reference; and GEMF-derived shared and full score representations. This section presents two benchmark analyses. The first compares methods across a common grid of low-dimensional representation sizes. The second uses reconstruction-based inner cross-validation and the one-standard-error rule to select representation dimension before evaluating external validity.

### 4.1. Dimension-matched benchmarking of compact subject-level signatures

We first compared methods across fixed low-dimensional representation sizes. For each rank *r* ∈ {1, 2, …, 12}, each PCA-based representation used the leading *r* principal components estimated within the training fold. The same outer cross-validation splits were used for all methods: 5 folds repeated 5 times, yielding 25 held-out evaluations. GEMF (shared) with shared rank *R* = *r* provides a “dimension-matched” comparison to rank-*r* PCA representations. GEMF (full) is shown as a secondary comparison because it uses *r* + 3 downstream scores rather than *r*. Figure 1 summarizes Δ*R*^2^ as a function of representation dimension for the outcomes age and LPS-1.

**Figure 1:**
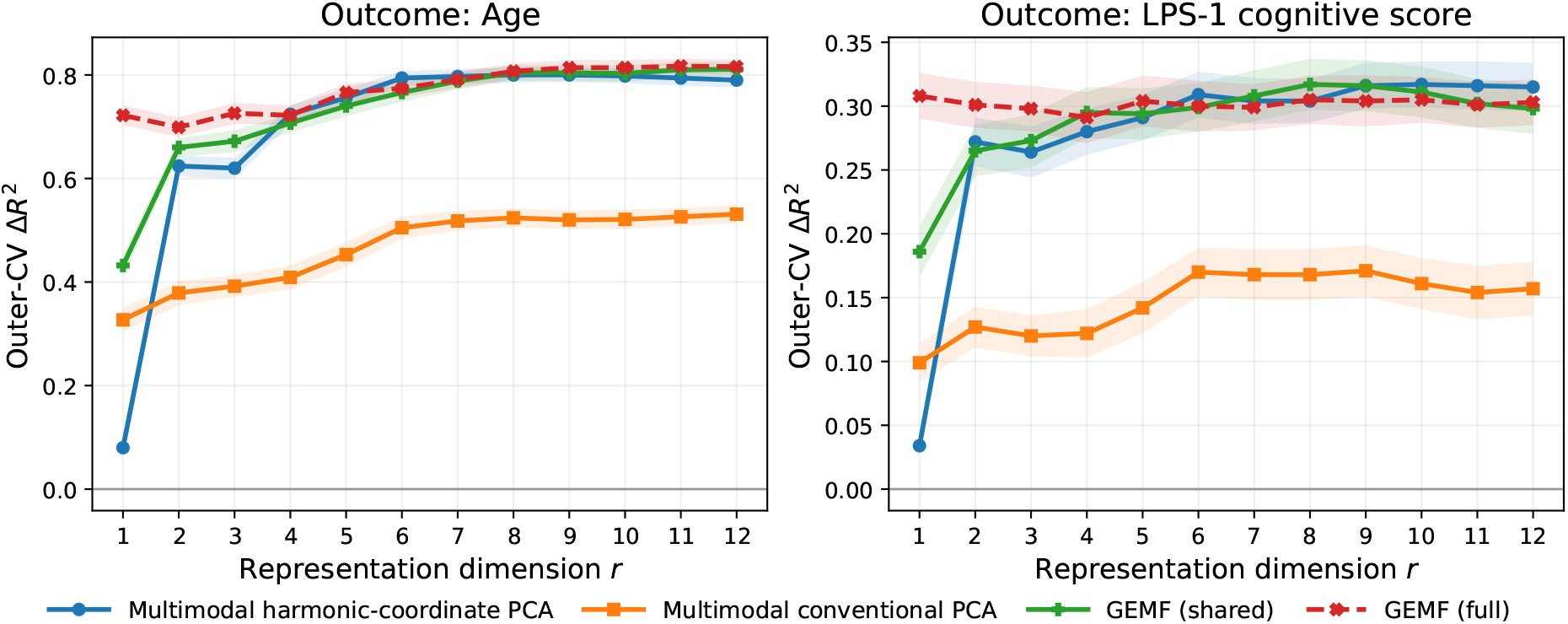
Dimension-matched benchmark of compact subject-level representations for external validation outcomes. Each curve shows the mean outer-cross-validated increase in *R*^2^, Δ*R*^2^, relative to a sex-only baseline model, across 25 held-out test folds. The *x*-axis gives the representation dimension *r*. For multimodal harmonic PCA and conventional PCA, *r* is the number of principal components. For GEMF (shared), *r* is the shared rank *R*. GEMF (full) is shown as a secondary reference; for shared rank *R* = *r*, it uses the *R*-dimensional shared score together with one modality-specific score for each modality, giving downstream dimension *r* + 3. Multimodal harmonic PCA and GEMF provide compact multimodal representations that substantially outperform conventional PCA for both age and LPS-1, with multimodal harmonic PCA reaching near-peak age prediction by moderate dimension.

For age (left panel), harmonic-coordinate representations substantially outperformed conventional PCA across the rank range in Figure 1. Multimodal conventional PCA improved gradually with increasing rank but reached only Δ*R*^2^ ≈ 0.531 at rank 12. In contrast, multimodal harmonic PCA was among the strongest approaches while requiring only moderate dimension: it reached Δ*R*^2^ ≈0.794 by rank 6 and plateaued around Δ*R*^2^ ≈ 0.800 by ranks 8–9. GEMF also performed strongly, with GEMF (shared) reaching Δ*R*^2^ ≈ 0.811 and GEMF (full) reaching Δ*R*^2^ ≈ 0.817 at rank 12. Thus, the best compact age signatures in the main comparison were based on cortical harmonic coordinates.

For the secondary LPS-1 outcome (right panel), the overall prediction signal was weaker than for age, but the same qualitative pattern was observed. Multimodal conventional PCA remained comparatively weak, peaking around Δ*R*^2^ ≈0.17. By contrast, multimodal harmonic PCA reached Δ*R*^2^ ≈ 0.317 around rank 10, and GEMF (shared) was also competitive, reaching Δ*R*^2^ ≈ 0.317 around rank 8. These results suggest that cortical harmonic coordinates capture subject-level variation that generalizes not only to chronological age but also to an independent cognitive measure.

The same comparison including all unimodal harmonic PCA representations is shown in Supplementary Figure S1. Those results show that the advantage of harmonic-coordinate representations was not driven solely by multimodal fusion: sMRI and rs-fMRI unimodal harmonic-coordinate PCA were themselves strong predictors for both age and LPS-1, while EEG unimodal harmonic-coordinate PCA was more modest but generally comparable to or better than conventional PCA.

Overall, the dimension-matched benchmark supports two conclusions. First, cortical harmonic coordinates provide strong compact subject-level brain signatures: unimodal harmonic summaries and multimodal harmonic PCA consistently outperformed conventional PCA. Second, multimodal integration in a common harmonic coordinate system is effective, not only for simple multimodal PCA-based fusion, but also for structured multiview factorization. GEMF matched or exceeded the strongest compact multimodal representations in this analysis while preserving modality-native structure and yielding shared factors that can be interpreted through sMRI, rs-fMRI, and EEG-specific loading objects.

### 4.2. Reconstruction-based rank selection

We next evaluated whether the same conclusions held when representation dimension was selected using training-set reconstruction rather than fixed (i.e., rather than “dimension-matched”) in advance. For each low-dimensional representation family, rank was selected from *r* = 1, …, 12 using the inner-cross-validation procedure described in Section 2.6. PCA-based methods used the normalized reconstruction error criterion in Equation 4, while GEMF used the criterion in Equation 5. For each outer split, the selected representation was refit on the full outer-training set and evaluated on the held-out subjects.

The selected ranks using one-standard-error rank selection were informative about the compactness of the representations. Multimodal conventional PCA selected a median rank of 6. Multimodal harmonic PCA selected a median rank of 8 (median across 25 outer test folds), whereas unimodal harmonic PCA representations selected median ranks near 11. GEMF selected a lower-dimensional shared representation, with median and modal selected rank 5. Thus, under the same reconstruction-based rule, GEMF favored a compact shared representation, while unimodal harmonic PCA generally required somewhat larger dimensionality.

After this rank selection, the main predictive conclusions using the median selected rank remained unchanged (Table 2). For age, multimodal harmonic PCA was the strongest selected-rank representation, with Δ*R*^2^ ≈ 0.798. sMRI harmonic PCA was close behind, with Δ≈ *R*^2^ 0.779. GEMF also remained strong: GEMF (full) achieved Δ*R*^2^ ≈ 0.767, while GEMF (shared) achieved Δ*R*^2^ ≈ 0.752. Conventional PCA was substantially weaker, with Δ*R*^2^ ≈ 0.513. The high-dimensional conventional ridge reference was competitive for age, with Δ*R*^2^ ≈ 0.744, but unlike the PCA and GEMF approaches it does not provide a compact subject-level representation.

**Table 2:** Reconstruction-based rank selection using inner cross-validation and the one-standard-error rule. Rank was selected from *r* = 1, …, 12 using normalized reconstruction error within each outer training set, without using the external outcomes. Performance is summarized over 25 outer test folds. Δ*R*^2^ is the increase in cross-validated *R*^2^ relative to a sex-only baseline model; Corr. is the prediction– outcome correlation; RMSE is the root mean squared prediction error. For GEMF, the selected rank is the shared rank *R*; GEMF (full) additionally includes one modality-specific score per modality.

| Representation | Median selected rank | Outcome: Age |  |  | Outcome: LPS-1 |  |  |
| --- | --- | --- | --- | --- | --- | --- | --- |
| | | $\Delta R^2$ | Corr. | RMSE | $\Delta R^2$ | Corr. | RMSE |
| Multimodal harmonic PCA | 8 | 0.798 | 0.901 | 8.799 | 0.308 | 0.557 | 3.425 |
| Multimodal conventional PCA | 6 | 0.513 | 0.731 | 13.493 | 0.164 | 0.418 | 3.740 |
| GEMF (shared) | 5 | 0.752 | 0.867 | 9.605 | 0.291 | 0.537 | 3.461 |
| GEMF (full) | 5 | 0.767 | 0.877 | 9.328 | 0.298 | 0.544 | 3.446 |
| sMRI harmonic PCA | 11 | 0.779 | 0.886 | 9.228 | 0.318 | 0.566 | 3.396 |
| fMRI harmonic PCA | 11 | 0.720 | 0.857 | 10.306 | 0.300 | 0.553 | 3.442 |
| EEG harmonic PCA | 11 | 0.511 | 0.727 | 13.385 | 0.177 | 0.427 | 3.717 |
| Multimodal conventional ridge | – | 0.744 | 0.864 | 9.921 | 0.189 | 0.476 | 3.661 |

For LPS-1, harmonic-coordinate methods again remained the strongest group. sMRI harmonic PCA achieved the highest selected-rank performance, with Δ*R*^2^ ≈0.318. Multimodal harmonic PCA achieved Δ*R*^2^ ≈ 0.308, and rs-fMRI harmonic PCA achieved Δ*R*^2^ ≈0.300. GEMF remained competitive, with Δ*R*^2^ ≈0.298 for GEMF (full) and Δ*R*^2^ ≈ 0.291 for GEMF (shared). In contrast, selected-rank multimodal conventional PCA achieved only Δ*R*^2^ ≈ 0.164. These results indicate that the advantage of harmonic-coordinate representations is not dependent on a particular prespecified rank; it persists when representation dimension is selected by an outcome-independent and training-set reconstruction criterion.

Taken together, the rank-selection analysis leads to a similar conclusion as the dimension-matched benchmark. Multimodal harmonic-coordiate PCA provides a compact and externally informative subject-level representation, selecting a lower median rank than unimodal harmonic PCA while retaining strong performance. The GEMF method selects an even lower median shared rank and remains predictively competitive, indicating that the structured multiview decomposition preserves useful subject-level variation while providing a more interpretable shared-factor representation. We next examine these GEMF shared factors directly to characterize how multiview structure is expressed across cortical spatial patterns, fMRI harmonic-coordinate loadings, EEG frequency and condition profiles, and subject-level age variation.

### 4.3. GEMF shared factors provide interpretable multiview signatures

The benchmark analyses above show that GEMF provides compact subject-level representations that remain competitive with PCA-based harmonic summaries. We next examined whether the fitted GEMF factors provide interpretable multiview structure. For this visualization analysis, we fit GEMF to the full MPI–LEMON common cohort using the median selected shared rank from the reconstruction-based rank-selection analysis, *R* = 5, with (*L*_*s*_, *L*_*f*_, *L*_*e*_) = (1, 1, 1). Because the sign of each factor is arbitrary, factor signs were oriented consistently for visualization; this sign convention does not affect the fitted reconstruction or downstream prediction.

For each shared factor *r*, GEMF yields a subject-level score *z*_*ir*_ and one loading object in each modality (Equation 1). The sMRI loading matrix **A**_*r*_ describes how the factor is expressed across harmonic modes and morphometric features. For a given morphometric feature *j* (e.g., cortical thickness), the corresponding cortical loading map can be reconstructed as

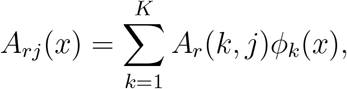

where *φ*_*k*_(*x*) is the *k*th cortical harmonic basis function evaluated at cortical location *x*. The rs-fMRI loading matrix **M**_*r*_ describes how the same shared factor loads on covariance entries between pairs of harmonic modes. Because **M**_*r*_ is a covariance-loading matrix rather than a scalar cortical field, we summarize its dominant cortical expression by projecting the leading eigenmode direction of **M**_*r*_ back to the cortical surface. Specifically, if **v**_*r*_ is the eigenvector of **M**_*r*_ associated with the eigenvalue of largest absolute magnitude, we define

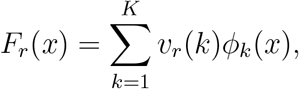

which is interpreted as the cortical harmonic pattern whose covariance contribution is most strongly represented in the factor-specific fMRI loading matrix. The EEG loading is decomposed into a spatial loading **a**_*r*_, frequency loading **b**_*r*_, and condition loading **c**_*r*_, allowing the same factor to be interpreted in terms of cortical spatial pattern, spectral profile, and eyes-open/eyes-closed contrast.

Figure 2 shows the representative shared factor *z*_5_, which had the strongest sex-adjusted association with age among the five GEMF shared factors (Figure 3). The cortical maps in Panels A–C show that this factor is spatially distributed rather than localized to a single region. The sMRI thickness loading in Panel A and the fMRI cortical pattern in Panel B in Figure 2 both show broad signed cortical gradients across lateral and medial cortex, indicating that the same subject-level factor is expressed in both morphology and harmonic covariance structure. The EEG cortical loading in Panel C shows a distinct but distributed cortical pattern in the forward-projected harmonic EEG representation. Thus, the shared factor does not impose identical spatial maps across modalities; instead, the same subject-level score is linked to modality-specific cortical loading patterns.

**Figure 2:**
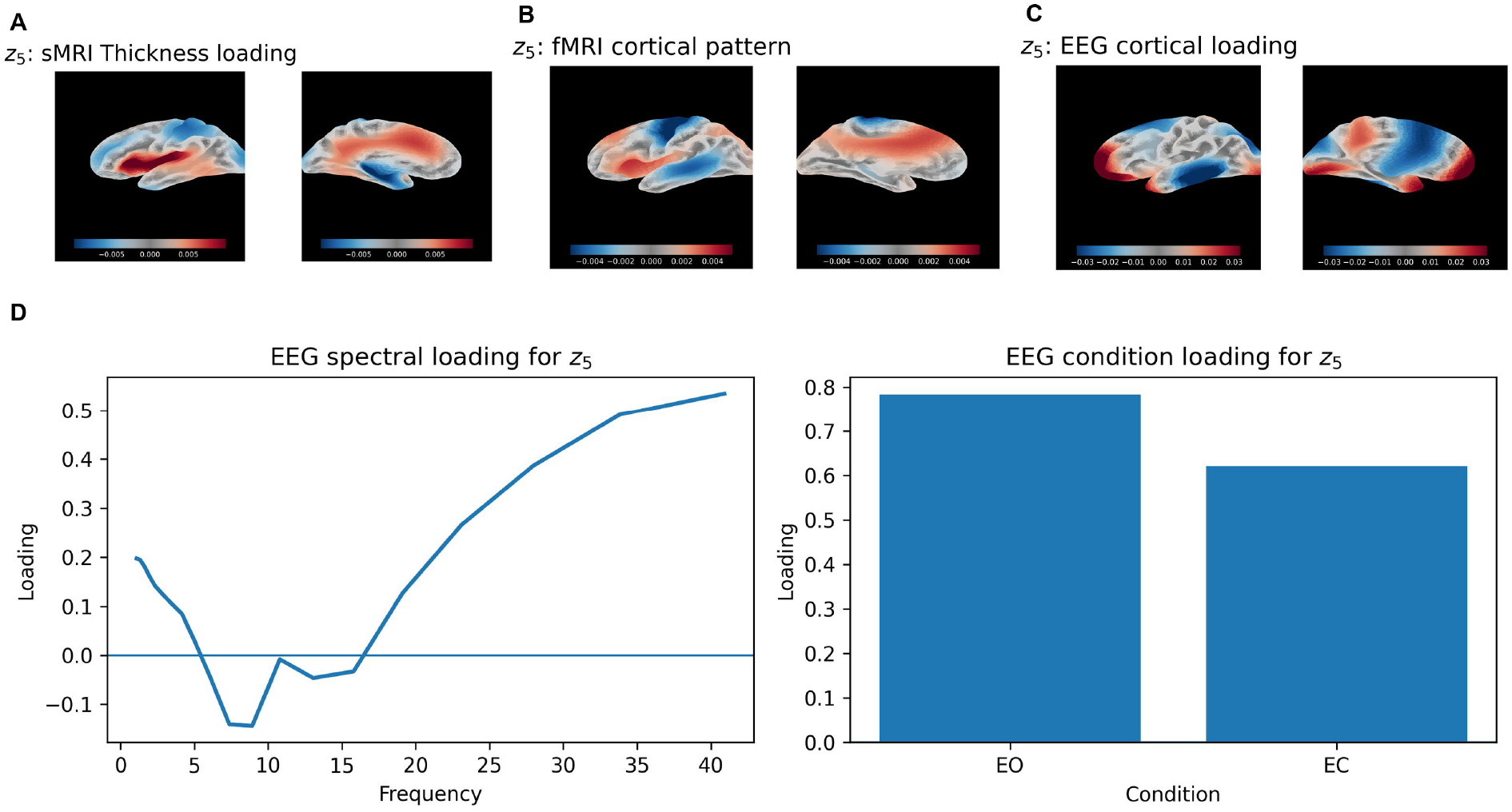
Representative GEMF shared factor *z*_5_, selected as the factor with the strongest sex-adjusted association with age. Panels show the corresponding sMRI thickness loading (A), fMRI cortical pattern, EEG cortical loading (B), and EEG spectral and condition loadings (C). The factor links a compact shared subject score to modality-specific loading objects that preserve the matrix, covariance, and tensor structure of the original modalities.

**Figure 3:**
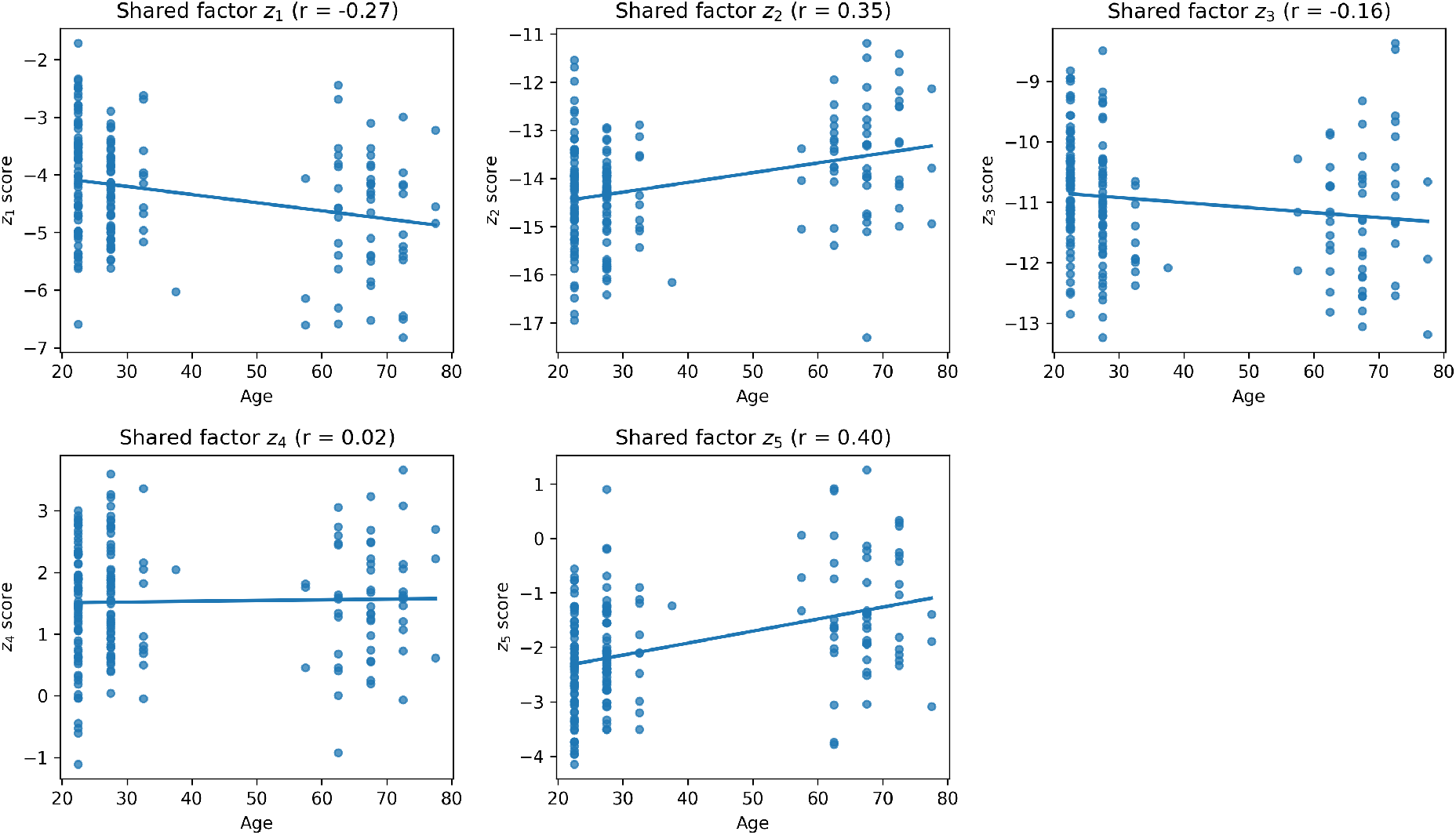
Associations between GEMF shared scores and chronological age. Each panel shows age on the horizontal axis and one inferred shared factor score on the vertical axis. These plots identify which shared multiview factors carry age-related variation.

The EEG loading profiles in Panel D provide an additional interpretation of the electrophysiological component of this factor. Since GEMF is fit to the log-power harmonic EEG tensor, the EEG loading factors are not constrained to be nonnegative. Under the chosen sign orientation, the EEG spectral loading is negative over a mid-frequency range and becomes increasingly positive at higher frequencies. The condition loading is positive for both eyes-open and eyes-closed rest, with stronger loading for eyes-open. These loadings should be interpreted as a spectral and condition contrast in the log-power representation associated with the factor, rather than as absolute increases or decreases in raw EEG power. Together, the cortical maps and loading profiles suggest that *z*_5_ captures an age-related multiview pattern expressed through cortical morphology, rs-fMRI harmonic covariance, and EEG spectral organization.

We also related the inferred shared factor scores to chronological age. In Figure 3, age was plotted on the horizontal axis and the shared score *z*_*ir*_ on the vertical axis for each factor, with sex-adjusted associations used as descriptive summaries. These plots show that age-related information is concentrated more strongly in some shared factors than others, motivating the detailed visualization of *z*_5_ in Figure 2. This illustrates the main interpretive advantage of GEMF: each shared factor consists of both a subject-level score and a set of modality-specific loading objects. Unlike a principal component score from a concatenated feature vector, whose loading vector mixes modalities and feature types in a single coordinate, a GEMF factor can be inspected separately as a cortical morphometric pattern, an rs-fMRI harmonic covariance pattern, and an EEG mode–frequency–condition profile. Thus, GEMF complements the predictive benchmarking results by providing a low-dimensional representation that is not only externally informative but also directly inspectable across modalities and cortical spatial scales.

## 5. Discussion

The present study evaluated whether cortical Laplace–Beltrami eigenmode coordinates can provide a common, interpretable, and practically useful subject-level representation across sMRI, rs-fMRI, and EEG. The results support three main conclusions. First, cortical harmonic coordinates produce compact subject-level signatures with strong external validity. Second, simple multimodal fusion in this coordinate system is feasible and effective. Third, GEMF provides a structured multiview decomposition that remains predictively competitive while adding interpretability through shared and modality-specific factors.

### Cortical harmonic coordinates provide compact subject-level signatures

Across the MPI– LEMON application, harmonic-coordinate representations consistently outperformed conventional low-dimensional atlas/sensor-based PCA representations. This was most pronounced for chronological age, where sMRI harmonic-coordinate PCA, rs-fMRI harmonic-coordinate PCA, multimodal harmonic-coordinate PCA, and GEMF all produced strong out-of-sample prediction. The advantage was not limited to age: the same broad pattern was observed for LPS-1, a secondary cognitive outcome, although the overall prediction signal was weaker. These findings suggest that cortical harmonic coordinates capture meaningful inter-individual variation rather than only compressing modality-specific data.

A central feature of the representation is its ordering by cortical spatial scale. Unlike conventional atlas-based summaries, which depend on a fixed parcellation, the LB basis provides a multiscale coordinate system in which low-order modes capture broad cortical organization and higher-order modes capture progressively finer spatial variation. This makes the resulting subject-level features interpretable not only as statistical summaries, but also as coordinates of cortical spatial organization. The strong performance of the unimodal harmonic-coordinate representations indicates that this geometry-aligned coordinate system is informative within individual modalities, while the multimodal results show that the same coordinate system can support cross-modal integration.

### Multimodal harmonic-coordinate PCA provides a strong and simple fusion benchmark

The multimodal harmonic-coordinate PCA representation was among the strongest-performing approaches. In the dimension-matched benchmark, it reached near-peak age prediction at moderate dimension and substantially outperformed conventional PCA. Under reconstruction-based rank selection, multimodal harmonic-coordinate PCA selected a median rank of 8 and retained strong performance for both age and LPS-1. This result is important because multimodal harmonic-coordinate PCA is methodologically simple: after each modality is expressed in harmonic coordinates, the modality-specific feature blocks are standardized, concatenated, and reduced by PCA. Its performance therefore supports the core representational claim of the paper: once sMRI, rs-fMRI, and EEG are expressed in a common geometry-aligned language, even simple linear fusion can yield compact and externally informative subject-level signatures.

At the same time, the supplementary unimodal comparisons show that the multimodal result was not driven solely by combining weak modality-specific summaries. sMRI and rsfMRI harmonic-coordinate PCA were themselves strong predictors, especially for age and LPS-1. EEG harmonic-coordinate PCA was more modest, but still generally comparable to or better than conventional PCA. Thus, the multimodal harmonic-coordinate representation combines informative modality-specific signals within a single low-dimensional harmonic coordinate system, rather than relying on one modality alone.

### GEMF adds structured interpretability while remaining competitive

GEMF was introduced as a structured multiview factorization that preserves the native organization of each modality while estimating shared and modality-specific subject scores. In contrast to simple concatenated PCA, GEMF models sMRI as a matrix of harmonic coefficients across morphometric fields, rs-fMRI as a harmonic covariance object, and EEG as a mode–frequency–condition tensor. This design allows each shared factor to be interpreted through modality-specific loading objects: cortical morphometric patterns, fMRI harmonic covariance structure, EEG cortical loadings, EEG spectral profiles, and condition effects.

The empirical results indicate that GEMF provides interpretability without losing the main predictive signal. In the dimension-matched benchmark, GEMF was highly competitive, particularly for age, and in some fixed-rank comparisons its shared or full score representation approached or exceeded multimodal harmonic-coordinate PCA. Under reconstruction-based rank selection, GEMF selected a lower median shared rank than multimodal harmonic-coordinate PCA and remained competitive for both age and LPS-1. This suggests that GEMF can preserve meaningful subject-level variation while providing a more structured and inspectable representation. The visualization analyses further show how shared GEMF factors can be examined across cortical spatial patterns, fMRI harmonic loadings, EEG frequency and condition profiles, and subject-level age variation. The simulation results provide additional support for using GEMF as a decomposition model. Under known data-generating mechanisms, GEMF reliably recovered shared subject-level structure across sample sizes and scenarios. Modality-specific recovery was strongest when modality-specific signal was substantial and more variable when modality-specific structure was weak relative to noise. These findings align with the intended in-terpretation of GEMF: the shared component is the most stable inferential target, while modality-specific factors provide useful but more signal-dependent structure.

### Implications for multimodal neuroimaging

The results suggest a practical strategy for multimodal neuroimaging studies. Rather than beginning with modality-specific atlases, sensor summaries, or separate dimension reductions, one can first express each cortical measurement in a common harmonic coordinate system. This creates a shared spatial language in which individual modalities can be analyzed separately, fused by simple methods such as PCA, or modeled jointly using structured decompositions such as GEMF. The present work focuses on cortical surface representations, but the broader idea is not restricted to the cortex in principle. Similar geometry-aligned representations could be developed for other anatomically meaningful surfaces or domains, provided that the relevant measurements can be represented as fields, time-varying fields, covariance objects, or structured tensors on a common manifold.

This point is especially relevant for future developmental, aging, and clinical neuroimaging studies. Subject-level signatures are most useful when they are compact, interpretable, and comparable across modalities and datasets (Arbabshirani et al., 2017; Marquand et al., 2019; Rutherford et al., 2022). Cortical harmonic coordinates provide a natural way to summarize distributed cortical organization while retaining information about spatial scale. In settings where some modalities are expensive, noisy, or missing, a common coordinate system may also help develop surrogate measures, for example EEG-only signatures that are informed by MRI-derived structure or fMRI-derived functional organization.

### Limitations

Several limitations should be noted. First, the empirical application used MPI–LEMON, a healthy adult cohort with a broad age range. Chronological age provides a strong external validation target, but it is not the same as disease prediction or clinical prognosis. The LPS-1 analysis provides a secondary cognitive validation outcome, but additional behavioral and clinical outcomes are needed to establish broader utility.

Second, the present GEMF implementation used a common *K* = 20 harmonic representation across modalities, focusing on coarse-to-intermediate spatial organization. This choice is well motivated by the spatial bandwidth shared across sMRI, rs-fMRI, and EEG, but future work should investigate multiresolution extensions that allow modality-specific bandwidths while maintaining shared cross-modal structure.

Third, the EEG representation depends on a forward-projected harmonic dictionary based on a template forward model. This provides a practical way to express scalp EEG in cortical harmonic coordinates, but subject-specific anatomy, head conductivity, sensor montage, and source-model uncertainty may affect the estimated EEG harmonic coefficients (Nunez and Srinivasan, 2006; Michel and Brunet, 2019). Future work should evaluate subject-specific forward models and uncertainty-aware EEG harmonic projection.

Fourth, GEMF remains a fixed-rank model with rank selected by reconstruction-based cross-validation. The one-standard-error rule provided a conservative and outcome-independent rank-selection strategy, but alternative approaches may be useful, including Bayesian shrinkage, stability-based selection, or prediction-constrained reconstruction criteria. We deliberately avoided selecting rank based on the external outcomes to preserve the validity of the benchmark, but future applications may require rank-selection strategies tailored to specific scientific goals.

Finally, while GEMF improves interpretability by preserving matrix, covariance, and tensor structure, it is more computationally involved than multimodal harmonic-coordinate PCA. The present results suggest that simple harmonic-coordinate PCA is an important reference method: it is easy to implement, performs strongly, and provides a useful benchmark against which more structured models should be judged. Future structured multiview models should therefore be evaluated not only by predictive performance, but also by whether they provide additional interpretability, stability, and scientific insight that justify their added complexity.

### Future directions

Several extensions are natural. First, the harmonic framework can be extended to age-varying or developmentally adaptive cortical bases, where the cortical geometry itself changes over time. This is particularly important for infancy and early childhood, where rapid cortical growth and folding may make fixed adult templates inappropriate. Second, longitudinal versions of GEMF could model within-subject trajectories of harmonic representations, separating stable subject-level differences from developmental or disease-related change. Third, multimodal harmonic-coordinate representations could be used to derive scalable surrogate biomarkers, such as EEG-only signatures informed by MRI and fMRI structure. Fourth, future work could incorporate additional modalities, including diffusion MRI, task fMRI, MEG, PET, or cortical maps of molecular and genetic organization, as long as they can be represented on the cortical surface or summarized in the harmonic domain.

## 6. Conclusion

Cortical Laplace–Beltrami eigenmode coordinates provide a practical common language for multimodal neuroimaging. In MPI–LEMON, harmonic-coordinate representations yielded compact subject-level signatures with strong external validity for age and a secondary cognitive outcome, outperforming conventional low-dimensional atlas/sensor-based PCA representations. Multimodal harmonic-coordinate PCA demonstrated that sMRI, rs-fMRI, and EEG can be fused effectively once represented in a shared geometry-aligned coordinate system. GEMF further showed that this common coordinate system can support structured multiview decomposition into shared and modality-specific components, preserving competitive prediction while enabling interpretable factor visualization across cortical scale, fMRI covariance structure, and EEG spectral/condition profiles.

Together, these findings support cortical harmonic coordinates as more than a compression tool. They provide a compact, interpretable, and multimodally aligned foundation for subject-level brain signatures, with potential applications in developmental neuroscience, aging research, clinical neuroimaging, and scalable multimodal biomarker development.

## Ethics

All analyses used de-identified human data from the publicly available MPI–LEMON dataset (Babayan et al., 2019). Ethical approval and informed consent procedures for participant recruitment and data collection were obtained by the original study investigators; the present work involves secondary analysis of these anonymized data.

## Data and Code Availability

The MPI–LEMON EEG and MRI data analyzed in this study are publicly available (see Babayan et al., 2019 for access and dataset details).

## Acknowledgements

The authors thank the MPI–LEMON investigators for making the dataset publicly available. Computations were performed on NYU’s High Performance Computing (HPC) resources, including the BigPurple cluster.

## Supplementary Material

Supplementary material is included in this submission PDF.

## Supplementary Materials

### S1. Supplementary Methods: Harmonic-Domain Feature Construction

This section describes the construction of the harmonic-domain sMRI, rs-fMRI, and EEG representations used in the main analyses. All three modalities were represented in a common cortical Laplace–Beltrami (LB) basis on the `fsaverage5` surface, using the same left-then-right hemisphere vertex ordering across modalities. The `fsaverage` template and surface registration tools were provided by FreeSurfer (Fischl, 2012), and the base MRI preprocessing used fMRIPrep (Esteban et al., 2019).

#### S1.1. Structural MRI morphometric fields

Morphometric processing was carried out for the primary structural MRI session using the subject-specific FreeSurfer reconstructions generated during the base preprocessing pipeline. For each subject and hemisphere, we extracted four canonical FreeSurfer surface measures: cortical thickness, sulcal depth, mean curvature, and surface area. These vertex-wise measures were treated as scalar fields on the cortical surface.

For each subject, hemisphere, and morphometric measure, the native-space scalar field was resampled to the `fsaverage5` template surface using FreeSurfer surface-based registration. The left- and right-hemisphere template fields were concatenated in a fixed vertex order, with all left-hemisphere vertices followed by all right-hemisphere vertices. This ordering matched the ordering used for the LB basis and for the rs-fMRI surface representation.

Let 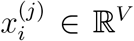 denote the concatenated `fsaverage5`-space scalar field for subject *I* and morphometric measure *j*, where *V* is the total number of template vertices across both hemispheres. Let Φ ∈ ℝ^*V* ×*K*^ denote the LB basis evaluated on the same template surface, and let **A**_*v*_ denote the diagonal matrix of vertex-area quadrature weights. The LB coefficient vector 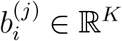 was obtained by the area-weighted least-squares projection

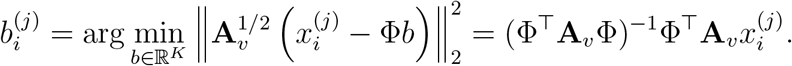

The full morphometry projection was computed using a larger set of LB coefficients, and the first *K*_*s*_ = 50 coefficients per measure were used for the main sMRI harmonic PCA analyses. For GEMF, the sMRI object was further restricted to the first *K* = 20 modes to match the common spatial dimension used across modalities.

The resulting structural harmonic object for subject *i* was the coefficient matrix

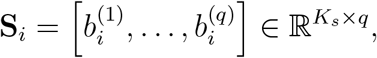

with *q* = 4 morphometric measures. Feature-standardized coefficients 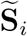 were constructed as described in the main Methods.

#### S1.2. Resting-state fMRI harmonic covariance

Resting-state fMRI data were preprocessed using fMRIPrep and represented on the cortical surface. After confound regression and temporal filtering, the surface-space BOLD signal at each time point was projected to the same LB basis used for the morphometric fields. Let *y*_*i*_(*t*) ∈ ℝ^*V*^ denote the surface BOLD field for subject *i* at time point *t*. The harmonic coefficient vector 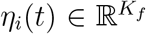 was obtained by the same area-weighted projection:

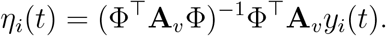

The main rs-fMRI harmonic PCA analyses used *K*_*f*_ = 50 harmonic coefficient time series. Subject-level rs-fMRI structure was summarized by the covariance matrix

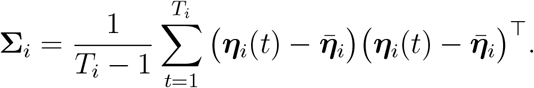

where 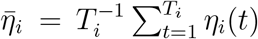. For modeling and feature construction, we used the log-Euclidean transform

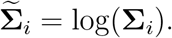

For GEMF, the rs-fMRI harmonic covariance object was restricted to the first *K* = 20 modes.

#### S1.3. EEG harmonic projection

Resting-state EEG was represented in the harmonic domain using a forward-projected cortical harmonic dictionary. Let 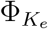 denote the first *K*_*e*_ cortical LB modes and let **L** ∈ ℝ^*M* ×*V*^ denote the template EEG forward operator mapping cortical surface activity to *M* scalp sensors. The sensor-space harmonic dictionary was

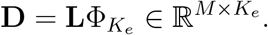

In this analysis, *K*_*e*_ = 20 and the EEG data contained *C* = 2 resting conditions, eyes-open and eyes-closed. For subject *i*, frequency bin *f*, and condition *c*, let **y**_*i*_(*f, c*) ∈ ℝ^*M*^ denote the log spectral-power topography across sensors. The harmonic coefficient vector was estimated by least squares:

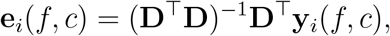

with optional ridge stabilization when needed for numerical conditioning. The coefficients were then arranged into the tensor

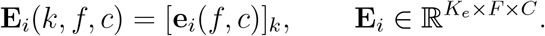

The main EEG representation retained *K*_*e*_ = 20 harmonic modes, *F* = 20 frequency bins, and *C* = 2 resting conditions.

#### S1.4. Analysis-ready harmonic representations

For each subject, the preprocessing pipeline produced three analysis-ready harmonic objects: the sMRI coefficient matrix 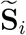, the log-Euclidean rs-fMRI harmonic covari-ance matrix 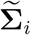, and the EEG mode–frequency–condition tensor **E**_*i*_. These objects were used directly to construct the unimodal harmonic PCA, multimodal harmonic PCA, and GEMF representations described in the main Methods.

### S2. Supplementary Methods: Computational Details for GEMF

#### S2.1. Fixed-rank GEMF objective

GEMF is estimated under fixed ranks (*R, L*_*s*_, *L*_*f*_, *L*_*e*_). For subject *i* = 1, …, *n*, the model is

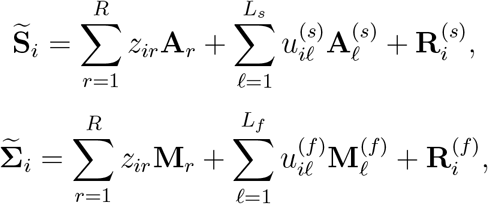

and

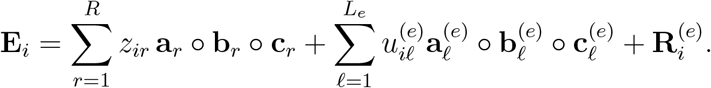

Here **z**_*i*_ = (*z*_*i*1_, …, *z*_*iR*_)^⊤^ is the shared score vector, while 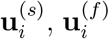 and 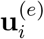 are modality-specific score vectors. To simplify notation in this supplement, modality-specific loading objects are denoted by superscripts (*s*), (*f*), (*e*).

The estimator minimizes the penalized objective

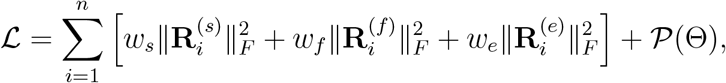

where Θ denotes all loading parameters. The shared-component penalty is

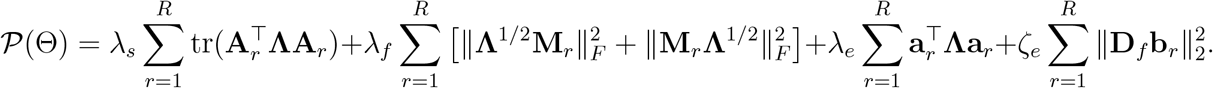

Here **Λ** = diag(*λ*_1_, …, *λ*_*K*_) contains the LB eigenvalues and **D**_*f*_ is a second-difference matrix over ordered frequency bins. In the MPI–LEMON application, we used *w*_*s*_ = *w*_*f*_ = *w*_*e*_ = 1, *λ*_*s*_ = *λ*_*f*_ = *λ*_*e*_ = 0.01, *ζ*_*e*_ = 0.01, a score ridge constant of 0.1, and weak ridge stabilization of 10^−6^ for modality-specific loading updates.

#### S2.2. Block-coordinate estimation

The fixed-rank GEMF estimator is computed by block-coordinate descent. Given current loading objects, subject scores are updated by penalized least squares. Given current scores, the sMRI and rs-fMRI loading objects are updated by penalized matrix regression, and EEG tensor factors are updated by alternating least squares for a CP-style rank-one tensor representation (Kolda and Bader, 2009).

##### S2.2.1. Shared score update

Let

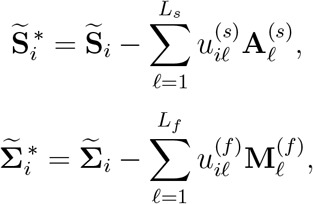

And

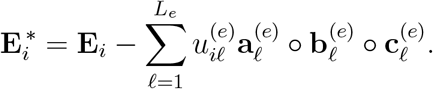

With all loading objects fixed, the shared score vector is updated as

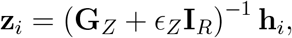

where *ϵ*_*Z*_ = 0.1 in the MPI–LEMON application. The matrix **G**_*Z*_ is a Gram matrix of the shared loading objects, with entries given by inner products among the shared components:

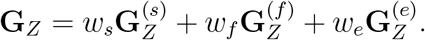

Specifically,

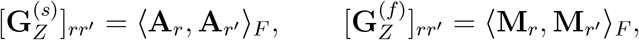

and

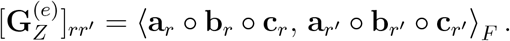

The vector **h**_*i*_ = (*h*_*i*1_, …, *h*_*iR*_)^⊤^ is the right-hand side of the penalized least-squares normal equations for **z**_*i*_. Its *r*th entry is

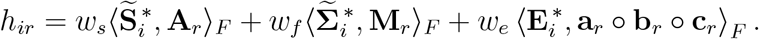

##### S2.2.2. Modality-specific score updates

The modality-specific score updates are analogous but use only the residual for the corresponding modality after removing the current shared component. For example, the sMRI-specific score vector is updated as

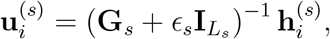

where

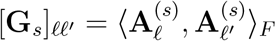

and

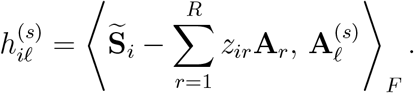

For rs-fMRI, the same update is applied using the modality-specific matrices 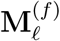. For EEG, the modality-specific loading for component 𝓁 is the rank-one tensor 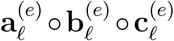, and the score update uses its Frobenius inner product with the EEG residual tensor.

##### S2.2.3. sMRI loading updates

For the *r*th shared sMRI loading matrix, define the current residual with all terms except *z*_*ir*_**A**_*r*_ removed:

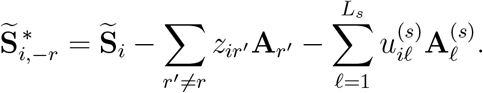

The update for **A**_*r*_ solves

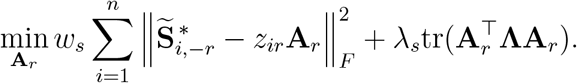

This yields

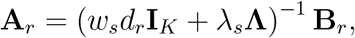

where 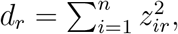, and 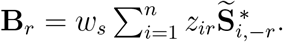. The sMRI-specific loading matrices 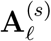 are updated in the same form but with weak ridge regularization rather than eigenvalue-weighted regularization. We used generic ridge regularization for modality-specific loadings because these terms serve primarily as nuisance components that absorb residual within-modality structure; the biologically interpretable scale-aware regularization is applied to the shared components, which are the primary object of inference.

##### S2.2.4. rs-fMRI loading updates

For the *r*th shared rs-fMRI loading matrix, define

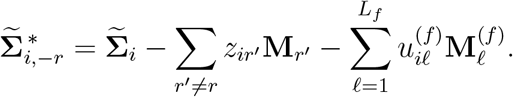

The update for **M**_*r*_ solves

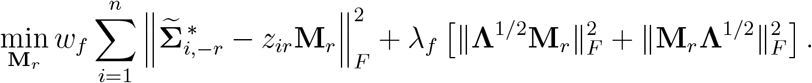

The elementwise update is 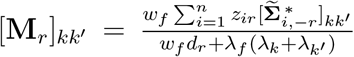, where *λ*_*k*_ and 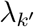 are the LB eigenvalues associated with harmonic modes *k* and *k*^′^. The denominator therefore penalizes high-frequency covariance patterns along both harmonic-mode dimensions. Because 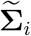 is symmetric, the fitted loading matrix is symmetrized after each update:

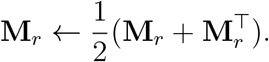

The rs-fMRI-specific loading matrices 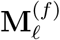 are updated analogously, using weak ridge regularization and the same symmetrization step.

##### S2.2.5. EEG tensor-factor updates

For the *r*th shared EEG component, define the current tensor residual with all terms except the *r*th shared EEG component removed:

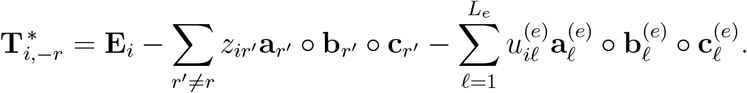

The shared EEG loading is represented as a rank-one tensor **a**_*r*_ ◦ **b**_*r*_ ◦ **c**_*r*_, where **a**_*r*_ is the spatial-mode loading, **b**_*r*_ is the frequency loading, and **c**_*r*_ is the condition loading. These factors are updated one at a time while holding the other two fixed.

Given (**b**_*r*_, **c**_*r*_), the spatial loading update is

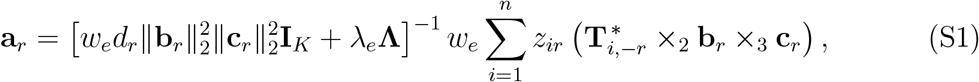

where 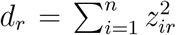, and ×_*m*_ denotes “contraction” of a tensor along mode *m* (Kolda and Bader, 2009; Sidiropoulos et al., 2017). In this three-way EEG tensor, 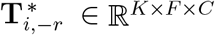 has modes corresponding to harmonic mode, frequency, and condition. Thus,the contraction 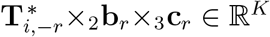 is a vector whose *k*th entry is 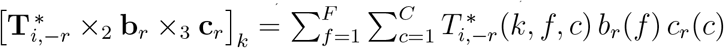. In other words, this operation collapses the frequency and condition modes using the current frequency and condition loadings, leaving a vector over cortical harmonic modes. The scalar multiplier 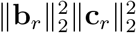 arises from the least-squares normal equations for the rank-one tensor factor **a**_*r*_ ◦**b**_*r*_ ◦**c**_*r*_. The parameter *λ*_*e*_ in (S1) controls eigenvalue-weighted shrinkage of the EEG spatial-mode loading, favoring smoother low-order cortical harmonic patterns unless higher-order modes are supported by the data.

Given (**a**_*r*_, **c**_*r*_), the frequency-profile update is

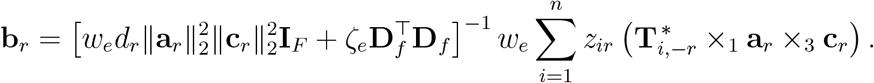

Here *ζ*_*e*_ (which we set to 0.01 in this paper) controls smoothness of the frequency loading through the second-difference penalty 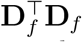.

Given (**a**_*r*_, **b**_*r*_), the condition-profile update is

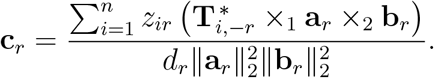

After each EEG factor update, the factorization is rescaled so that

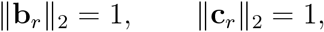

with the corresponding amplitude absorbed into **a**_*r*_. The modality-specific EEG factors are updated using the same alternating least-squares structure but with weak ridge regularization and without the shared spatial or frequency smoothness penalties. As with the sMRI and rs-fMRI modality-specific components, these terms are included to absorb residual modality-specific variation rather than to serve as the primary interpretable shared structure.

#### S2.3. Initialization, normalization, and convergence

The implementation uses a deterministic warm-start strategy. First, a concatenated subject-by-feature matrix is formed from the vectorized harmonic objects vec 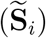, vech 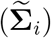, and vec(**E**_*i*_), after applying modality weights. The first *R* left singular vectors of this matrix initialize the shared score matrix. Given these initial shared scores, the shared sMRI and rs-fMRI loading matrices are initialized by least-squares projection of the corresponding harmonic objects onto the initial scores. Each shared EEG component is initialized by a rank-one approximation to the score-weighted average EEG tensor. Modality-specific components are initialized from residuals after the initial shared fit.

After each score update, columns of the shared score matrix and each modality-specific score matrix are centered and rescaled to unit variance. The corresponding scaling is absorbed into the associated loading objects, so that fitted reconstructions are unchanged. This provides a practical identifiability convention for computation.

The algorithm stops when the relative change in the penalized objective satisfies

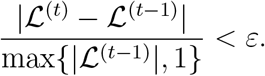

In the MPI–LEMON application, we used *ε* = 10^−5^ and a maximum of 100 block-coordinate iterations.

#### S2.4. Held-out score inference

In cross-validation, GEMF loadings are estimated using only the outer-training subjects. For held-out subjects, the fitted loading objects are kept fixed and subject scores are inferred by penalized least squares. Specifically, for a held-out subject with harmonic objects 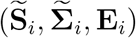, we solve for either the shared score **z**_*i*_ alone or the full score vector

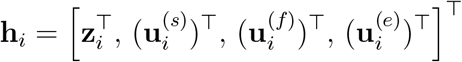

with all loading objects fixed. The same inference step is applied to training and held-out subjects before downstream prediction, ensuring that GEMF (shared) and GEMF (full) use consistent score definitions across cross-validation folds.

### S3. Supplementary Methods: Conventional Atlas- and Sensor-Based Feature Construction

This section describes the construction of “conventional” subject-level features used as comparison representations in the MPI–LEMON application. These features reflect commonly used atlas- and sensor-based summaries for sMRI, rs-fMRI, and EEG, in contrast to the cortical harmonic-coordinate representations used in the main analyses.

#### S3.1. Subject alignment

All conventional feature matrices were restricted to the same analytic subjects used for the harmonic-coordinate analyses. Only subjects with available harmonic representations, conventional feature summaries, age, sex, and the relevant secondary outcome were retained for a given analysis. For multimodal analyses, the sMRI, rs-fMRI, and EEG feature matrices were ordered using the same subject identifiers before feature concatenation.

#### S3.2. Conventional sMRI atlas features

For structural MRI, conventional morphometric features were obtained from FreeSurfer atlas summaries using the Destrieux cortical parcellation (Destrieux et al., 2010). The Destrieux atlas contains 74 cortical regions per hemisphere, yielding 148 bilateral cortical regions. For each subject, we extracted regional summaries for cortical thickness, surface area, and mean curvature from the FreeSurfer parcellation statistics. These regional values were concatenated across hemispheres and morphometric measures to form the conventional sMRI feature vector

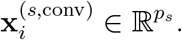

Here *p*_*s*_ denotes the number of retained regional morphometric features after merging the available atlas summaries.

#### S3.3. Conventional rs-fMRI atlas features

For resting-state fMRI, conventional features were based on atlas-level functional connectivity. Preprocessed BOLD data were represented on the cortical surface and summarized using the Destrieux cortical parcels. Let **r**_*i*_(*t*) ∈ ℝ^*P*^ denote the vector of parcel time series for subject *i*, where *P* is the number of retained Destrieux cortical parcels after excluding unavailable or empty parcels. Nuisance regression and temporal filtering were applied consistently with the rs-fMRI preprocessing used for harmonic feature construction (Esteban et al., 2019). We computed the parcel-level correlation matrix

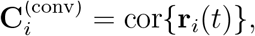

and vectorized its upper triangle:

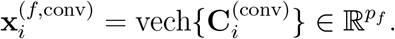

This vector provides a conventional atlas-based resting-state connectivity representation.

#### S3.4. Conventional EEG sensor features

For EEG, conventional resting-state features were computed in sensor space rather than in the cortical harmonic dictionary. Eyes-open and eyes-closed resting data were processed separately and then merged at the subject level. For each condition, we computed band-limited spectral summaries over theta, alpha, and beta frequency ranges. Sensor-level power summaries were aggregated into regional channel groups to reduce dimensionality and improve stability. We also included alpha peak frequency and condition-specific alpha-power summaries, which are commonly used resting-state EEG descriptors.

Let

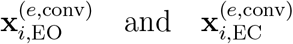

denote the conventional EEG feature vectors for eyes-open (EO) and eyes-closed (EC) conditions. The final conventional EEG feature vector was

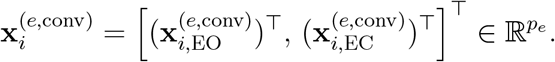

#### S3.5. Combined conventional multimodal representation

The conventional multimodal representation was formed by concatenating the conventional sMRI, rs-fMRI, and EEG feature vectors:

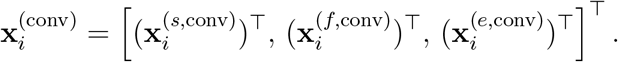

This representation was used to construct two comparison models.

First, the conventional PCA representation was obtained by standardizing all conventional features within each training fold and applying PCA to the concatenated conventional feature matrix. For a given rank *r*, the leading *r* principal component scores were used as the low-dimensional conventional subject representation. This provides a dimension-matched comparator to multimodal harmonic PCA.

Second, the conventional ridge representation used the full standardized conventional feature vector directly in ridge regression. The ridge tuning parameter was selected within the training data using inner cross-validation and then applied to the held-out outer-test subjects. This model was included as a high-dimensional conventional benchmark and was not constrained to match the dimension of the low-rank harmonic or PCA representations.

#### S3.6. Preprocessing within cross-validation

All standardization, PCA estimation, ridge tuning, and predictive modeling steps were performed within the training folds of the outer cross-validation. For each outer split, feature means and standard deviations were estimated using training subjects only and applied to held-out test subjects. PCA loadings for the conventional PCA baseline were estimated using training subjects only, and test-subject principal component scores were obtained by projection onto the training-derived PCA loadings. Ridge regression models were fit using standardized training features and then applied to standardized held-out test features. This procedure ensured that no information from held-out subjects entered feature normalization, dimension reduction, tuning, or model fitting.

### S4. Supplementary Methods and Results: Simulation Study

#### S4.1. Simulation data-generating mechanism

The simulation study evaluated GEMF under controlled data-generating mechanisms in which the true shared and modality-specific components were known. For subject *i* = 1, …, *n*, data were generated directly in the same object spaces used by GEMF:

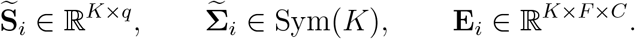

We fixed (*K, q, F, C*) = (20, 4, 16, 2) and true shared rank *R* = 3. In scenarios with modality-specific structure, we used (*L*_*s*_, *L*_*f*_, *L*_*e*_) = (1, 1, 1). Sample size was varied over *n* ∈ {50, 100, 200}, with 100 replications per setting.

The true shared score matrix **Z** ∈ ℝ^*n*×*R*^ was generated with approximately orthonormal columns. Modality-specific score matrices **U**_*s*_, **U**_*f*_, and **U**_*e*_ were generated similarly and orthogonalized against the shared score matrix. Loading objects were generated with spatial weights favoring lower-order harmonic modes. For sMRI and rs-fMRI, modality-specific loading objects were orthogonalized against the span of the corresponding shared loading objects in vectorized loading space. For EEG, modality-specific rank-one tensor loadings were orthogonalized against the span of the vectorized shared EEG tensor loadings and then projected back to a rank-one tensor approximation.

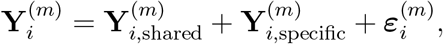

where *m* ∈ {*s, f, e*} denotes sMRI, rs-fMRI, or EEG. The three components were scaled so that their total squared Frobenius norms, summed across subjects, followed prespecified proportions within each modality. Specifically, for *c* ∈ {shared, specific, noise},

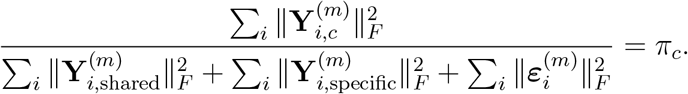

The same target proportions were used for sMRI, rs-fMRI, and EEG within each scenario. We considered three scenarios: shared-only, with proportions (0.70, 0.00, 0.30) for shared, modality-specific, and noise components; shared+specific, with proportions (0.45, 0.35, 0.20); and weak-specific, with proportions (0.35, 0.05, 0.60). The fixed-rank GEMF estimator was fit using the true ranks in each scenario. Shared-structure recovery is reported in the main text. Modality-specific recovery and rank-selection results are reported in Sections S4.2 and S4.3.

#### S4.2. Modality-specific recovery

In scenarios containing modality-specific structure, we evaluated recovery of the modality-specific score subspaces and modality-specific components. Score recovery was summarized by the mean canonical correlation between estimated and true modality-specific score subspaces. Component recovery was summarized by

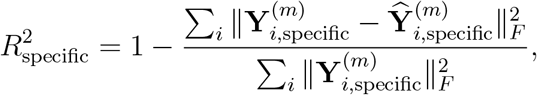

separately for each modality.

**Table S1:** Supplementary fixed-rank simulation recovery of modality-specific structure across sample sizes. Values are mean (SD) across 100 replications. Specific-score recovery is the mean canonical correlation between estimated and true modality-specific score subspaces. Specific reconstruction is 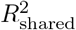 for the true modality-specific component. Entries are not applicable for the shared-only scenario.

| Scenario | $n$ | Specific-score recovery | | | Specific reconstruction $R^2$ | | |
| --- | --- | --- | --- | --- | --- | --- | --- |
|  |  | sMRI | fMRI | EEG | sMRI | fMRI | EEG |
| Shared+specific | 50 | 0.996 (0.001) | 0.998 (0.001) | 0.802 (0.396) | 0.960 (0.029) | 0.967 (0.026) | 0.696 (0.544) |
|  | 100 | 0.996 (0.001) | 0.999 (0.001) | 0.930 (0.252) | 0.977 (0.015) | 0.980 (0.015) | 0.889 (0.358) |
|  | 200 | 0.996 (0.001) | 0.999 (0.000) | 0.911 (0.283) | 0.985 (0.006) | 0.988 (0.010) | 0.870 (0.396) |
| Weak-specific | 50 | 0.920 (0.020) | 0.966 (0.008) | 0.816 (0.363) | 0.499 (0.072) | 0.644 (0.039) | 0.689 (0.562) |
|  | 100 | 0.925 (0.010) | 0.969 (0.004) | 0.762 (0.406) | 0.672 (0.040) | 0.793 (0.025) | 0.619 (0.692) |
|  | 200 | 0.930 (0.008) | 0.971 (0.003) | 0.896 (0.283) | 0.765 (0.024) | 0.866 (0.013) | 0.819 (0.498) |

Table S1 shows that modality-specific recovery was strong in the shared+specific scenario, especially for sMRI and rs-fMRI. EEG-specific recovery was more variable at smaller sample sizes, reflecting the greater difficulty of recovering a structured mode– frequency–condition tensor component. In the weak-specific scenario, modality-specific recovery was attenuated relative to the shared+specific scenario, especially for reconstruction of the weaker sMRI and EEG-specific components. These results complement the main simulation table by showing that modality-specific recovery is more sensitive to sample size and signal strength than recovery of the shared component.

#### S4.3. Cross-validated shared-rank selection

We also evaluated whether the shared rank *R* could be selected using the same reconstruction-based strategy used in the empirical analyses. Candidate ranks were *R* ∈ {1, …, 12}, with true rank *R* = 3. For each candidate rank, GEMF was fit within cross-validation folds and evaluated using modality-balanced normalized reconstruction error. We compared two criteria: the rank minimizing the mean cross-validated reconstruction error and the one-standard-error rule. The one-standard-error rule selected the smallest rank whose mean reconstruction error was within one standard error of the minimum.

Table S2 shows that choosing the rank by the raw mean cross-validation error tended to overselect, often favoring high ranks. In contrast, the one-standard-error rule selected ranks close to the true value. Exact recovery of the true rank was highest in the shared-only scenario and remained high in the weak-specific scenario. In the shared+specific scenario, the selected rank was often *R* = 3 or a neighboring rank, reflecting the greater difficulty of separating shared from modality-specific structure when both are present. These results support the use of the one-standard-error rule as a conservative, outcome-independent rank-selection strategy.

### S5. Supplementary Methods and Results: Application

#### S5.1. Supplementary dimension-matched benchmark including unimodal harmonic PCA

The main application figure emphasizes the compact multimodal representations: multimodal harmonic PCA, conventional PCA, GEMF (shared), and GEMF (full). To assess whether the performance of harmonic-coordinate representations was driven primarily by multimodal fusion or was already present within individual modalities, we also repeated the dimension-matched benchmark including the unimodal sMRI, rs-fMRI, and EEG harmonic PCA representations.

**Table S2:** Supplementary simulation results for cross-validated shared-rank selection. Candidate ranks were *R* = 1, …, 12, and the true shared rank was *R* = 3. The one-standard-error rule selects the smallest rank whose mean modality-balanced normalized reconstruction error is within one standard error of the minimum. Prop. exact denotes the proportion of replications in which the one-standard-error rule selected the true shared rank. Best by CV mean denotes the rank minimizing the mean modality-balanced normalized reconstruction error.

| Scenario | $n$ | True $R$ | One-SE rule | Prop. exact (One-SE) | Best by CV mean |
| --- | --- | --- | --- | --- | --- |
| Shared-only | 50 | 3 | 3.01 (0.10) | 0.99 | 11.74 (1.44) |
|  | 100 | 3 | 3.04 (0.24) | 0.97 | 10.49 (3.29) |
|  | 200 | 3 | 3.05 (0.22) | 0.95 | 10.31 (3.38) |
| Shared+specific | 50 | 3 | 3.32 (0.47) | 0.68 | 3.60 (1.35) |
|  | 100 | 3 | 3.41 (0.53) | 0.61 | 3.73 (1.47) |
|  | 200 | 3 | 3.43 (0.70) | 0.63 | 4.20 (2.53) |
| Weak-specific | 50 | 3 | 3.11 (0.49) | 0.94 | 11.86 (0.93) |
|  | 100 | 3 | 3.21 (0.78) | 0.86 | 11.90 (0.90) |
|  | 200 | 3 | 3.51 (1.37) | 0.79 | 11.90 (0.72) |

Supplementary Figure S1 shows that unimodal harmonic representations were themselves strongly informative. For chronological age, sMRI harmonic PCA and rs-fMRI harmonic PCA substantially outperformed conventional PCA across most ranks, while EEG harmonic PCA showed more modest performance. For LPS-1, sMRI and rs-fMRI harmonic PCA again provided strong subject-level signatures, with performance comparable to the multimodal harmonic PCA and GEMF representations at moderate ranks. These results indicate that the benefit of the harmonic-coordinate framework is not limited to cross-modal fusion: the common cortical harmonic coordinate system also yields informative modality-specific subject representations.

#### S5.2. Supplementary GEMF shared-factor visualizations

The main text displays the representative GEMF shared factor selected for detailed interpretation. To show the full set of fitted shared factors, Supplementary Figures S2–S5 display the remaining four GEMF shared factors using the same visualization format. Each figure links one shared subject-level score to its modality-specific loading objects: an sMRI cortical loading map, an rs-fMRI cortical pattern, an EEG cortical loading map, and EEG spectral and condition loadings. Supplementary Figure S6 shows all four sMRI morphometric loading maps for the representative factor shown in the main text.

**Figure S1:**
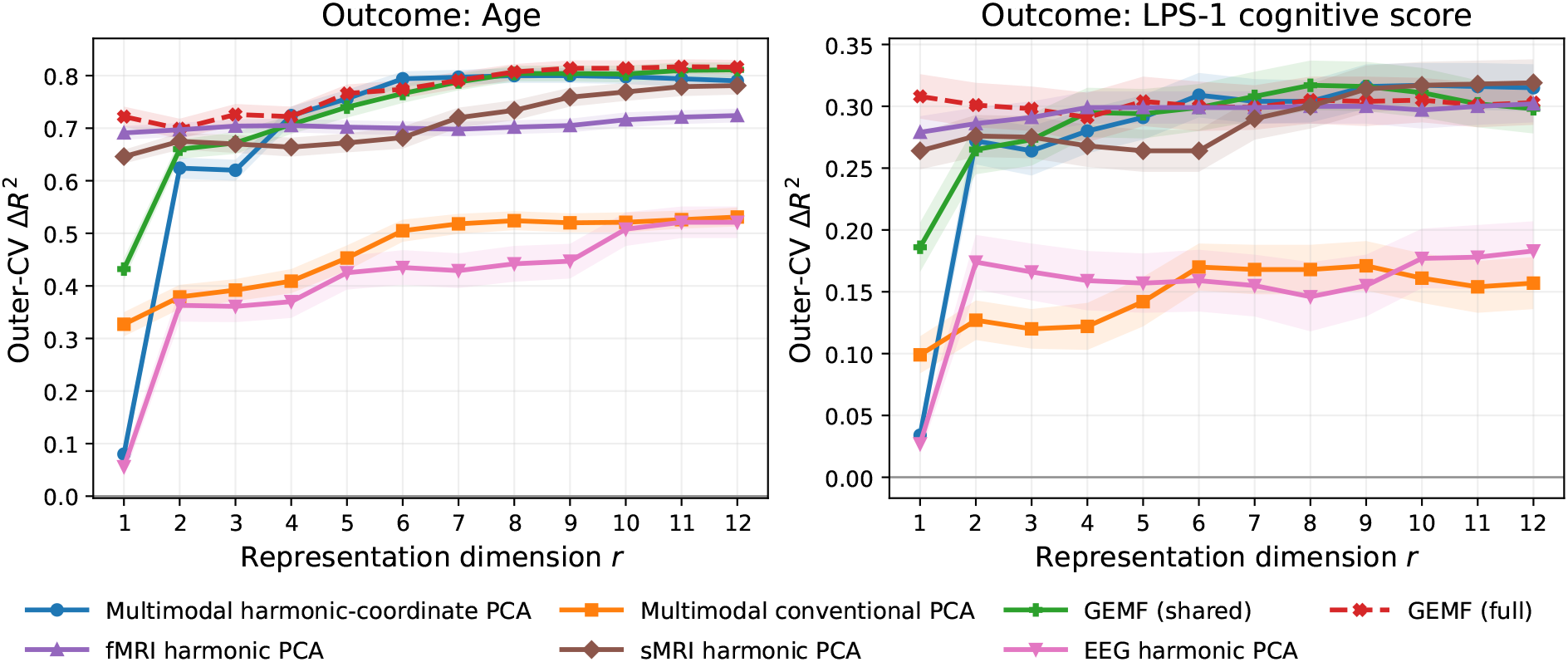
Supplementary dimension-matched benchmark including unimodal harmonic PCA representations. Each curve shows the mean outer-cross-validated Δ*R*^2^ relative to a sex-only baseline model across 25 held-out test folds. The *x*-axis gives the representation dimension *r*. Multimodal harmonic PCA and conventional PCA use *r* principal components. The unimodal harmonic PCA curves use *r* principal components from the corresponding modality-specific harmonic-coordinate feature block. GEMF (shared) uses the *R* = *r* shared scores, whereas GEMF (full) additionally includes one modality-specific score for each of sMRI, fMRI, and EEG. The supplementary comparison shows that the advantage of harmonic-coordinate representations over conventional PCA is not driven solely by multimodal fusion: both sMRI and fMRI harmonic PCA provide strong unimodal subject-level signatures, while multimodal harmonic PCA and GEMF remain competitive across outcomes.

**Figure S2:**
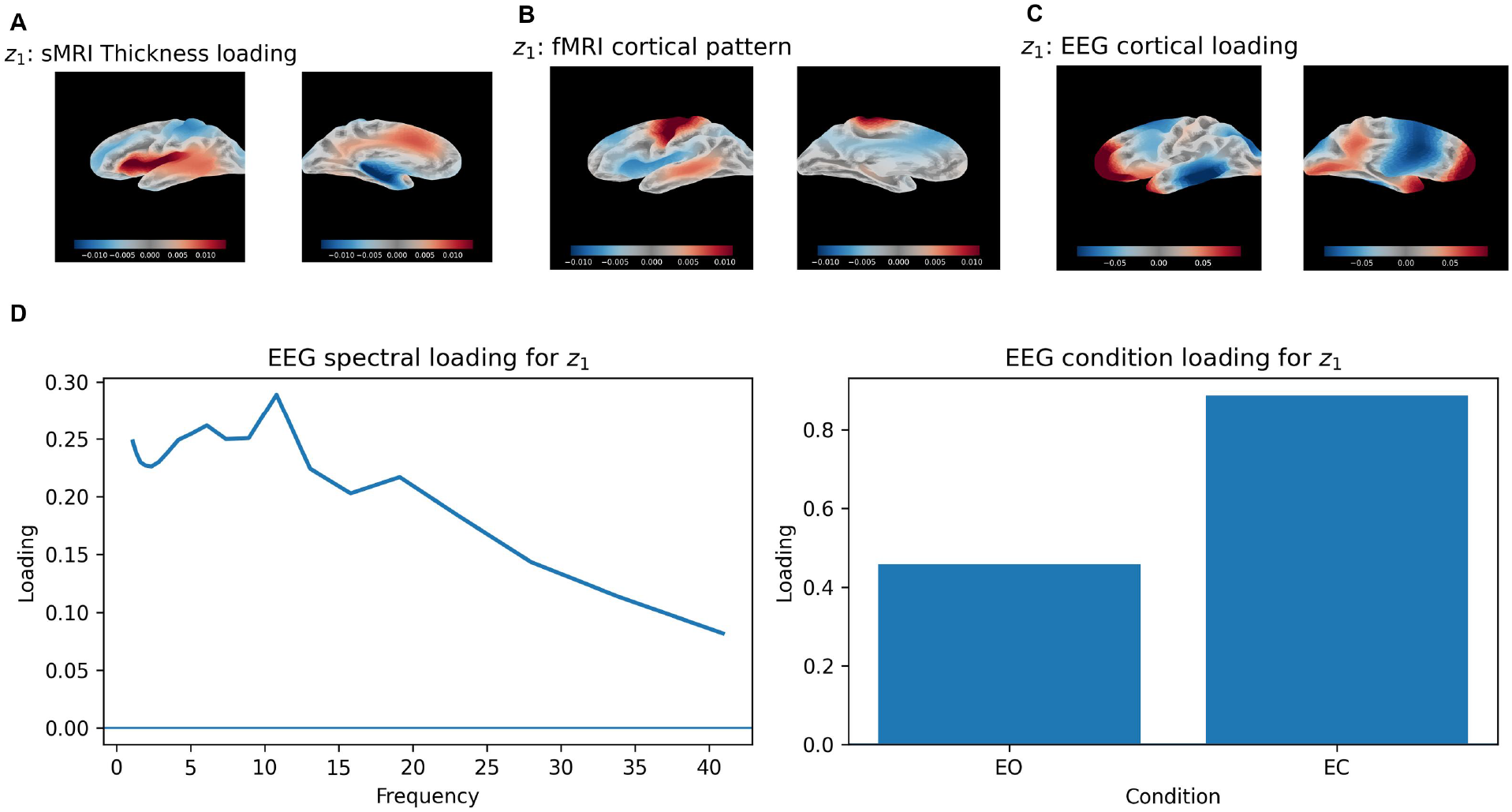
Supplementary visualization of GEMF shared factor *z*_1_. Panels show the modality-specific loading objects associated with the same shared subject-level factor: an sMRI cortical loading map, an rs-fMRI cortical pattern, an EEG cortical loading map, and EEG spectral and condition loadings. The visualization illustrates how a single shared factor is expressed across anatomy, hemodynamic covariance structure, and electrophysiological spectral/condition profiles.

**Figure S3:**
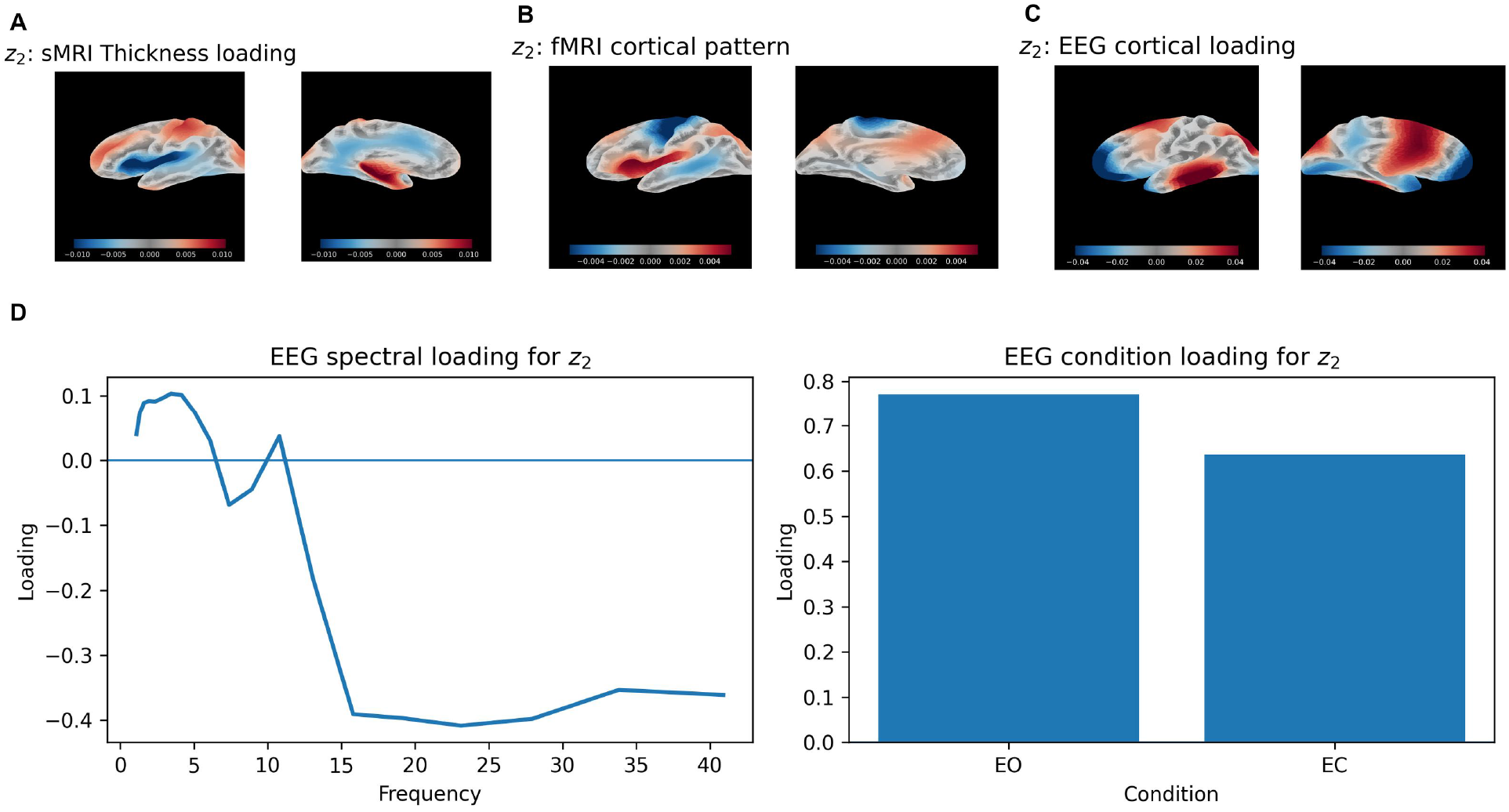
Supplementary visualization of GEMF shared factor *z*_2_. Panels show the modality-specific loading objects associated with the same shared subject-level factor: an sMRI cortical loading map, an rs-fMRI cortical pattern, an EEG cortical loading map, and EEG spectral and condition loadings. Together with the other GEMF factor visualizations, this figure shows that the shared representation is not a single undifferentiated score space, but a set of interpretable multiview components.

**Figure S4:**
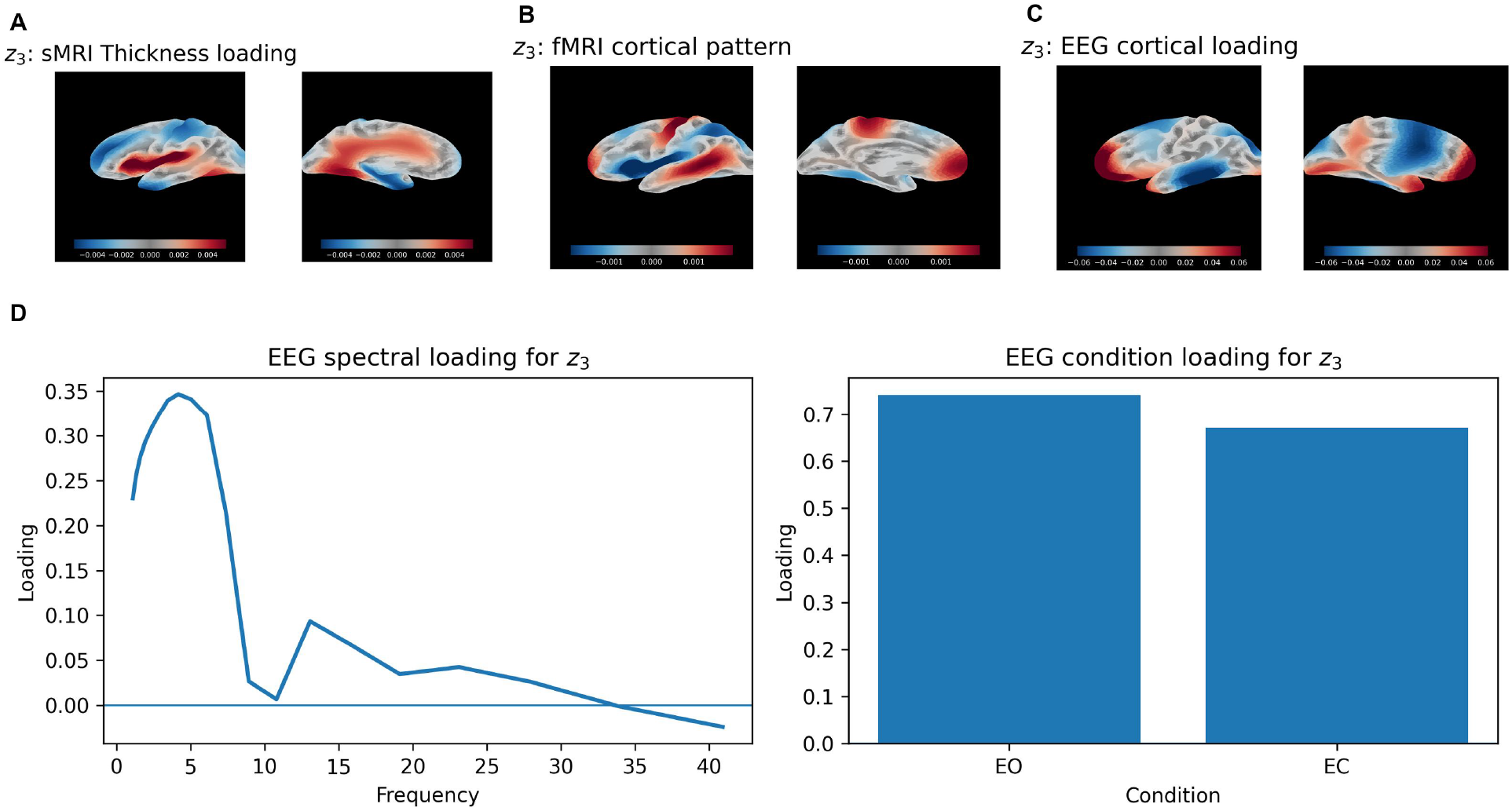
Supplementary visualization of GEMF shared factor *z*_3_. Panels show the modality-specific loading objects associated with the same shared subject-level factor: an sMRI cortical loading map, an rs-fMRI cortical pattern, an EEG cortical loading map, and EEG spectral and condition loadings. The factor-specific loadings retain the native structure of each modality while linking them through a common shared subject score.

**Figure S5:**
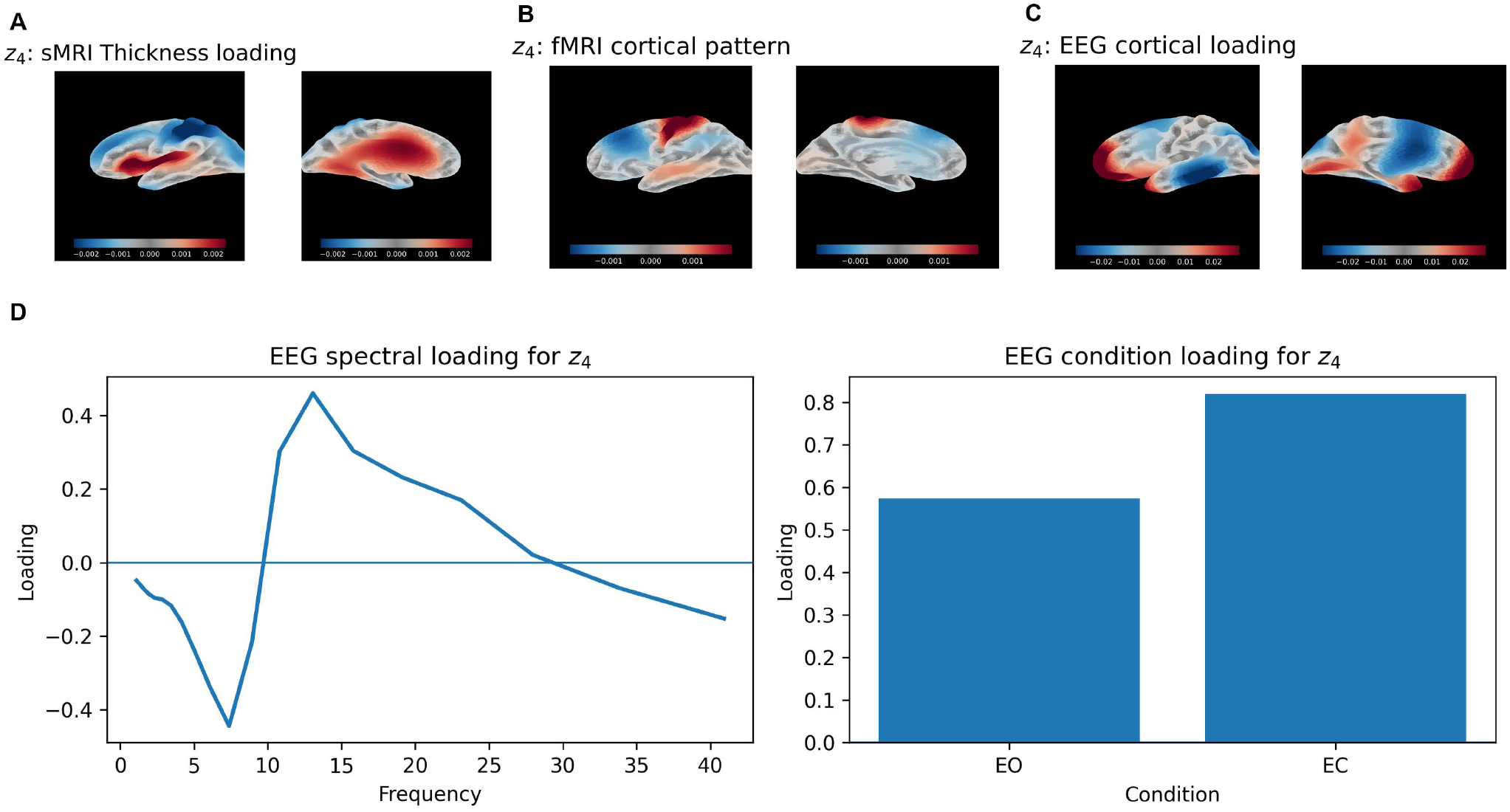
Supplementary visualization of GEMF shared factor *z*_4_. Panels show the modality-specific loading objects associated with the same shared subject-level factor: an sMRI cortical loading map, an rs-fMRI cortical pattern, an EEG cortical loading map, and EEG spectral and condition loadings. This display complements the main-text representative factor by showing another shared multiview component estimated from the same *R* = 5, *L*_*s*_ = *L*_*f*_ = *L*_*e*_ = 1 GEMF model.

**Figure S6:**
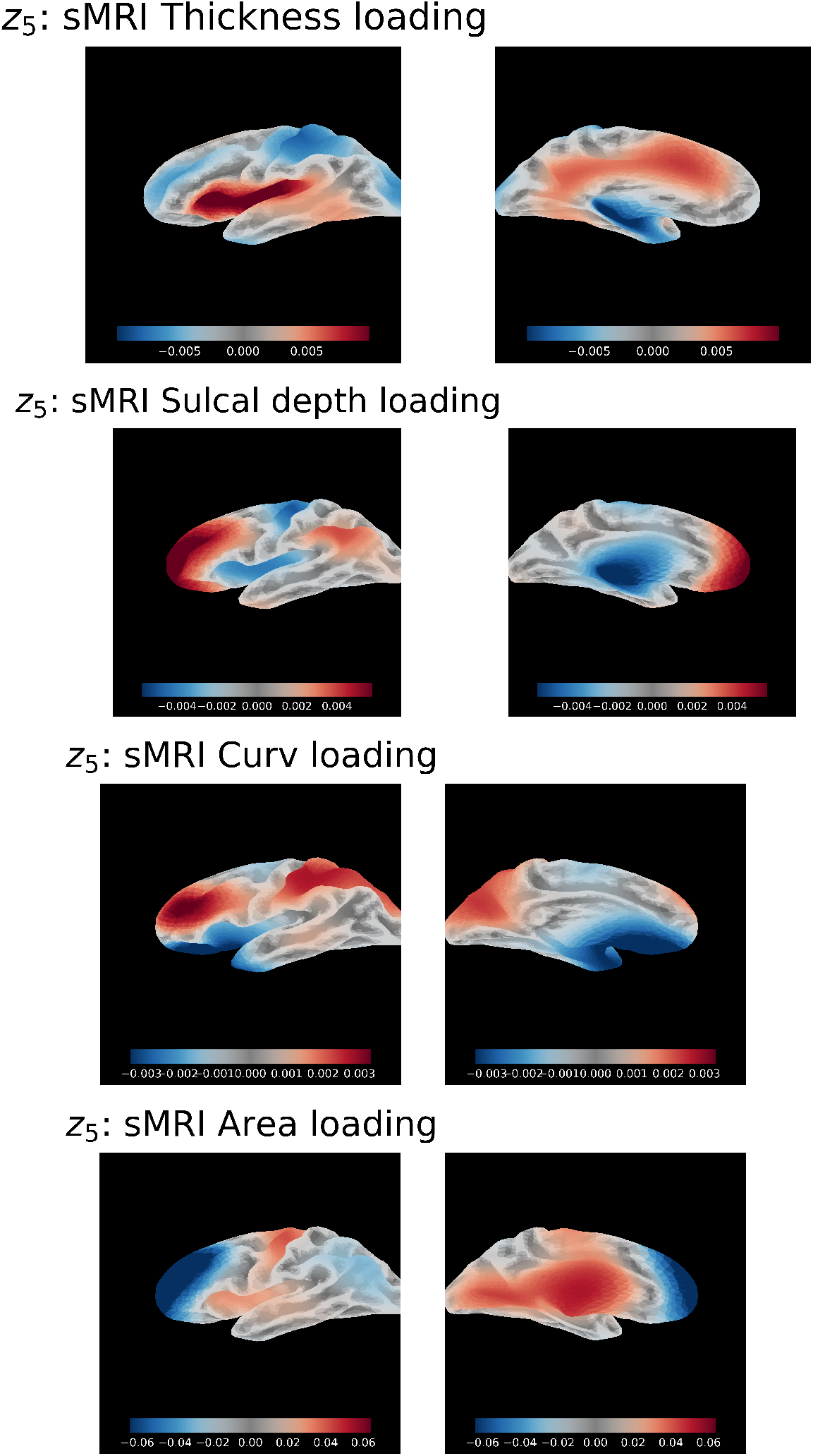
sMRI morphometric loading maps for GEMF shared factor *z*_5_. Panels show the cortical loading maps for the four morphometric fields included in the sMRI harmonic object. Displaying all morphometric loadings for the same shared factor illustrates how GEMF represents the anatomical component as a matrix over cortical harmonic mode and morphometric feature, rather than as a single vectorized loading.

## References

Arbabshirani, M. R., Plis, S., Sui, J., and Calhoun, V. D. (2017). Single subject prediction of brain disorders in neuroimaging: Promises and pitfalls. NeuroImage, 145:137–165.

Argelaguet, R., Arnol, D., Bredikhin, D., Deloro, Y., Velten, B., Marioni, J. C., and Stegle, O. (2020). MOFA+: a statistical framework for comprehensive integration of multi-modal single-cell data. Genome Biology, 21:111.

Argelaguet, R., Velten, B., Arnol, D., Dietrich, S., Zenz, T., Marioni, J. C., Buettner, F., Huber, W., and Stegle, O. (2018). Multi-omics factor analysis—a framework for unsupervised integration of multi-omics data sets. Molecular Systems Biology, 14(6):e8124.

Atasoy, S., Donnelly, I., and Pearson, J. (2016). Human brain networks function in connectome-specific harmonic waves. Nature Communications, 7:10340.

Babayan, A., Erbey, M., Kumral, D., Reinelt, J. D., Reiter, A. M. F., Röbbig, J., Schaare, H. L., Uhlig, M., Anwander, A., Bazin, P.-L., Horstmann, A., Lampe, L., Nikulin, V. V., Okon-Singer, H., Preusser, S., Pampel, A., Rohr, C. S., Sacher, J., Thöne-Otto, A., Trapp, S., Nierhaus, T., Altmann, D., Arelin, K., Blöchl, M., Bongartz, E., Breig, P., Cesnaite, E., Chen, S., Cozatl, R., Czerwonatis, S., Dambrauskaite, G., Dreyer, M., Enders, J., Engelhardt, M., Fischer, M. M., Forschack, N., Golchert, J., Golz, L., Guran, C. A., Hedrich, S., Hentschel, N., Hoffmann, D. I., Huntenburg, J. M., Jost, R., Kosatschek, A., Kunzendorf, S., Lammers, H., Lauckner, M. E., Mahjoory, K., Kanaan, A. S., Mendes, N., Menger, R., Morino, E., Näthe, K., Neubauer, J., Noyan, H., Oligschläger, S., Panczyszyn-Trzewik, P., Poehlchen, D., Putzke, N., Roski, C., Schaller, M.-C., Schieferbein, A., Schlaak, B., Schmidt, R., Gorgolewski, K. J., Schmidt, H. M., Schrimpf, A., Stasch, S., Voss, M., Wiedemann, A., Margulies, D. S., Gaebler, M., and Villringer, A. (2019). A mind-brain-body dataset of MRI, EEG, cognition, emotion, and peripheral physiology in young and old adults. Scientific Data, 6:180308.

Bi, X., Tang, X., Yuan, Y., Zhang, Y., and Qu, A. (2021). Tensors in statistics. Annual Review of Statistics and Its Application, 8:345–368.

Biswal, B. B., Mennes, M., Zuo, X.-N., Gohel, S., Kelly, C., Smith, S. M., Beckmann, C. F., Adelstein, J. S., Buckner, R. L., Colcombe, S., Dogonowski, A.-M., Ernst, M., Fair, D., Hampson, M., Hoptman, M. J., Hyde, J. S., Kiviniemi, V. J., Kötter, R., Li, S.-J., Lin, C.-P., Lowe, M. J., Mackay, C., Madden, D. J., Madsen, K. H., Margulies, D. S., Mayberg, H. S., McMahon, K., Monk, C. S., Mostofsky, S. H., Nagel, B. J., Pekar, J. J., Peltier, S. J., Petersen, S. E., Riedl, V., Rombouts, S. A. R. B., Rypma, B., Schlaggar, B. L., Schmidt, S., Seidler, R. D., Siegle, G. J., Sorg, C., Teng, G.-J., Veijola, J., Villringer, A., Walter, M., Wang, L., Weng, X., Whitfield-Gabrieli, S., Williamson, P., Windischberger, C., Zang, Y.-F., Zhang, H.-Y., Castellanos, F. X., and Milham, M. P. (2010). Toward discovery science of human brain function. Proceedings of the National Academy of Sciences of the United States of America, 107(10):4734–4739.

Calhoun, V. D. and Sui, J. (2016). Multimodal fusion of brain imaging data: A key to finding the missing link(s) in complex mental illness. Biological Psychiatry: Cognitive Neuroscience and Neuroimaging, 1(3):230–244.

Cao, T., Pang, J. C., Segal, A., Chen, Y.-C., Aquino, K. M., Breakspear, M., and Fornito, A. (2024). Mode-based morphometry: A multiscale approach to mapping human neuroanatomy. Human Brain Mapping, 45:e26640.

Cohen, M. X. (2014). Analyzing Neural Time Series Data: Theory and Practice. MIT Press, Cambridge, MA.

Destrieux, C., Fischl, B., Dale, A., and Halgren, E. (2010). Automatic parcellation of human cortical gyri and sulci using standard anatomical nomenclature. NeuroImage, 53(1):1–15.

Eilers, P. H. C. and Marx, B. D. (1996). Flexible smoothing with B-splines and penalties. Statistical Science, 11(2):89–121.

Esteban, O., Markiewicz, C. J., Blair, R. W., Moodie, C. A., Isik, A. I., Erramuzpe, A., Kent, J. D., Goncalves, M., DuPre, E., Snyder, M., Oya, H., Ghosh, S. S., Wright, J., Durnez, J., Poldrack, R. A., and Gorgolewski, K. J. (2019). fMRIPrep: A robust preprocessing pipeline for functional MRI. Nature Methods, 16(1):111–116.

Ferreira, F. S., Mihalik, A., Adams, R. A., Ashburner, J., and Mourao-Miranda, J. (2022). A hierarchical Bayesian model to find brain-behaviour associations in incomplete data sets. NeuroImage, 249:118854.

Fischl, B. (2012). Freesurfer. NeuroImage, 62(2):774–781.

Gholamipourbarogh, N., Eggert, E., Münchau, A., Frings, C., and Beste, C. (2024). EEG tensor decomposition delineates neurophysiological principles underlying conflict-modulated action restraint and action cancellation. NeuroImage, 295:120667.

Groves, A. R., Beckmann, C. F., Smith, S. M., and Woolrich, M. W. (2011). Linked independent component analysis for multimodal data fusion. NeuroImage, 54(3):2198–2217.

Hastie, T., Tibshirani, R., and Friedman, J. (2009). The Elements of Statistical Learning: Data Mining, Inference, and Prediction. Springer, New York, 2 edition.

Horn, W. (1983). Leistungsprüfsystem (LPS). Hogrefe, Göttingen.

Jorge, J., van der Zwaag, W., and Figueiredo, P. (2014). EEG–fMRI integration for the study of human brain function. NeuroImage, 102:24–34.

Klami, A., Virtanen, S., and Kaski, S. (2013). Bayesian canonical correlation analysis. Journal of Machine Learning Research, 14:965–1003.

Klami, A., Virtanen, S., Leppäaho, E., and Kaski, S. (2015). Group factor analysis. IEEE Transactions on Neural Networks and Learning Systems, 26(9):2136–2147.

Klimesch, W. (1999). EEG alpha and theta oscillations reflect cognitive and memory performance: a review and analysis. Brain Research Reviews, 29(2–3):169–195.

Kolda, T. G. and Bader, B. W. (2009). Tensor decompositions and applications. SIAM Review, 51(3):455–500.

Kucukboyaci, N. E., Kemmotsu, N., Leyden, K. M., Girard, H. M., Tecoma, E. S., Iragui, V. J., and McDonald, C. R. (2014). Integration of multimodal MRI data via PCA to explain language performance. NeuroImage: Clinical, 5:197–207.

Lock, E. F., Hoadley, K. A., Marron, J. S., and Nobel, A. B. (2013). Joint and individual variation explained (JIVE) for integrated analysis of multiple data types. The Annals of Applied Statistics, 7(1):523–542.

Luppi, A. I., Vohryzek, J., Kringelbach, M. L., Mediano, P. A. M., Craig, M. M., Adapa, R., Carhart-Harris, R. L., Roseman, L., Pappas, I., Peattie, A. R. D., Manktelow, A. E., Sahakian, B. J., Finoia, P., Williams, G. B., Allanson, J., Pickard, J. D., Menon, D. K., Atasoy, S., and Stamatakis, E. A. (2023). Distributed harmonic patterns of structure-function dependence orchestrate human consciousness. Communications Biology, 6:117.

Marquand, A. F., Kia, S. M., Zabihi, M., Wolfers, T., Buitelaar, J. K., and Beckmann, C. F. (2019). Conceptualizing mental disorders as deviations from normative functioning. Molecular Psychiatry, 24(10):1415–1424.

Michel, C. M. and Brunet, D. (2019). EEG source imaging: A practical review of the analysis steps. Frontiers in Neurology, 10:325.

Nunez, P. L. and Srinivasan, R. (2006). Electric Fields of the Brain: The Neurophysics of EEG. Oxford University Press, New York, 2 edition.

Pang, J. C., Aquino, K. M., Oldehinkel, M., Robinson, P. A., Fulcher, B. D., Breakspear, M., and Fornito, A. (2023). Geometric constraints on human brain function. Nature, 618(7965):566–574.

Park, H. G. (2026). Forward-projected cortical eigenmodes provide an efficient sensor-space representation of resting-state EEG. Brain Topography, 39:33.

Power, J. D., Cohen, A. L., Nelson, S. M., Wig, G. S., Barnes, K. A., Church, J. A., Vogel, A. C., Laumann, T. O., Miezin, F. M., Schlaggar, B. L., and Petersen, S. E. (2011). Functional network organization of the human brain. Neuron, 72(4):665–678.

Reuter, M., Wolter, F.-E., Shenton, M., and Niethammer, M. (2009). Laplace–Beltrami eigenvalues and topological features of eigenfunctions for statistical shape analysis. Computer-Aided Design, 41(10):739–755.

Ruppert, D., Wand, M. P., and Carroll, R. J. (2003). Semiparametric Regression. Number 12 in Cambridge Series in Statistical and Probabilistic Mathematics. Cambridge University Press, Cambridge.

Rutherford, S., Kia, S. M., Wolfers, T., Fraza, C., Zabihi, M., Dinga, R., Berthet, P., Worker, A., Verdi, S., Andrews, D. S., Han, L., Tso, I. F., Cardoner, N., Chaim-Avancini, T., Hoekstra, P. J., Mennes, M., Andreassen, O. A., Westlye, L. T., and Marquand, A. F. (2022). The normative modeling framework for computational psychiatry. Nature Protocols, 17(7):1711–1734.

Seo, S. and Chung, M. K. (2011). Laplace–beltrami eigenfunction expansion of cortical manifolds. In 2011 IEEE International Symposium on Biomedical Imaging: From Nano to Macro, pages 372–375. IEEE.

Shen, X., Finn, E. S., Scheinost, D., Rosenberg, M. D., Chun, M. M., Papademetris, X., and Constable, R. T. (2017). Using connectome-based predictive modeling to predict individual behavior from brain connectivity. Nature Protocols, 12(3):506–518.

Sidiropoulos, N. D., De Lathauwer, L., Fu, X., Huang, K., Papalexakis, E. E., and Faloutsos, C. (2017). Tensor decomposition for signal processing and machine learning. IEEE Transactions on Signal Processing, 65(13):3551–3582.

Sui, J., Adali, T., Yu, Q., Chen, J., and Calhoun, V. D. (2012). A review of multivariate methods for multimodal fusion of brain imaging data. Journal of Neuroscience Methods, 204(1):68–81.

Sui, J., He, H., Yu, Q., Chen, J., Rogers, J., Pearlson, G. D., Mayer, A. R., Bustillo, J., Canive, J. M., and Calhoun, V. D. (2013). Three-way (N-way) fusion of brain imaging data based on mCCA+jICA and its application to discriminating schizophrenia. NeuroImage, 66:119–132.

Sui, J., Jiang, R., Bustillo, J., and Calhoun, V. D. (2020). Neuroimaging-based individualized prediction of cognition and behavior for mental disorders and health: Methods and promises. Biological Psychiatry, 88(11):818–828.

Sui, J., Zhi, D., and Calhoun, V. D. (2023). Data-driven multimodal fusion: approaches and applications in psychiatric research. Psychoradiology, 3:kkad026.

Wingeier, B. M., Nunez, P. L., and Silberstein, R. B. (2001). Spherical harmonic decomposition applied to spatial-temporal analysis of human high-density electroencephalogram. Physical Review E, 64(5):051916.

Zhou, G., Cichocki, A., Zhang, Y., and Mandic, D. P. (2016). Group component analysis for multiblock data: Common and individual feature extraction. IEEE Transactions on Neural Networks and Learning Systems, 27(11):2426–2439.

